# A Novel Model of Pancreatic Cancer Dormancy Reveals Mechanistic Insights and a Dormancy Gene Signature with Human Relevance

**DOI:** 10.1101/2020.04.13.037374

**Authors:** Crissy Dudgeon, Chris R. Harris, Ying Chen, Bassel Ghaddar, Anchal Sharma, Mihir M. Shah, Arthur I. Roberts, Anthony Casabianca, Eric A. Collisson, Vinod P. Balachandran, Paula M. Vertino, Subhajyoti De, Darren R. Carpizo

## Abstract

Latent recurrences following curative-intent pancreatic cancer surgery is a major clinical problem thought to be due to the reactivation of dormant tumor cells that disseminate before the primary tumor has been removed. How dormancy is established and ultimately reversed to drive recurrence is poorly understood. Here we introduce a novel model of pancreatic cancer dormancy that mimics early and latent survival outcomes of resected patients. Using single-cell transcriptomics we compared primary, dormant, and reactivated tumor cells and found the primary and reactivated tumor cell transcriptomes clustered together with and away from the dormant tumor cells. Using a chromatin accessibility assay we found dormancy exhibits large scale changes in chromatin remodeling. Dormant tumor cells express cancer stem cell markers that are lost upon reactivation and are chemotherapy resistant. We identified a dormancy gene signature and investigated this in patients undergoing surgery for localized PC by isolating cells from the primary tumor and liver disseminated tumor cells (DTCs) for single-cell transcriptomics. We found the signature correlated with DTCs indicating that these cells are dormant at the time of surgery. The signature also identified CCL5 as a novel dormancy marker in PC. Mechanisms of PC dormancy include upregulation of the transcriptional repressor Dec2 which drives quiescence, monoallelic suppression of the mutant KRAS allele by DNA methylation, and immunoregulation. We conclude that PC dormancy is a highly plastic and heterogeneous cellular state governed by tumor cell autonomous and non-autonomous mechanisms.

**One Sentence Summary:** A novel model of resectable pancreatic cancer reveals pancreatic cancer dormancy is characterized by significant cellular plasticity, heterogeneity and chromatin remodeling

## Introduction

Latent metastatic recurrence following surgery with curative intent is one of the most significant clinical problems in oncology. These latent recurrences are due to tumor cells that have disseminated to distant organ sites that are occult at the time of surgery and remain dormant for a period of time (sometimes years) before reactivating to form tumors. The mechanisms of metastatic tumor cell dormancy are poorly understood primarily because of the inability to identify and capture such cells from patients. While various animal models of metastatic tumor cell dormancy have been reported (*1, 2*), none involve orthotopic primary tumor growth in immunocompetent animals followed by surgical resection with spontaneous metastatic dissemination. These models are challenging because they require survival surgery as well as extended periods of follow up time.

Pancreatic cancer (PC) is a highly lethal malignancy (projected second leading cause of cancer death in the United States by 2030) with a proclivity for metastatic dissemination (*3*) (*4*). In patients undergoing curative intent surgery for PC, recurrence is highly frequent (over 80%) and is most commonly due to disseminated disease rather than local recurrence. Moreover, both early and latent recurrent phenotypes are observed (*5*). There are currently no mouse models of pancreas cancer that reflect the biology of the resected patient. Genetically engineered mouse models (GEMMs) of PC are inadequate for the study of MTCD in that the disease pathology affects the entire gland and thus a model of late stage (stage IV) disease. Here we have developed the first mouse model of early stage resectable pancreatic cancer for the study of PC dormancy. This model provides a number of insights into the biology of pancreatic cancer dormancy and reveals a gene signature that has relevance to the study of dormancy in human PC patients.

## Results

### A novel model of early stage pancreatic cancer that recapitulates outcomes of resected patients

To gain insight into the mechanisms that contribute to pancreatic cancer cell dormancy and recurrence after surgical resection, we sought to develop a mouse model that recapitulates the patterns of recurrence and survival outcomes of resected patients. Ideally, such a model would exhibit spontaneous dissemination prior to surgical resection of the primary tumor as the physiology of surgery has been shown to impact recurrence and dormancy (*6*) and involve an immunocompetent host, since emerging data states that immunosurveillance is a strong mediator of dormancy (*7*) (*8*). To construct this model, we adapted a syngeneic orthotopic model using a cell line (Ink4a.1) derived from the Pdx-1-Cre; Kras^G12D/+^; p16^Ink4a -/-^ model that expresses luciferase and mCherry allowing for in vivo tumor monitoring and downstream lineage tracing (*9*). Cells were injected into the distal pancreas of FVB mice to form a primary tumor prior to undergoing a distal pancreatectomy/splenectomy, which is the standard surgery for distal pancreatic adenocarcinoma (Fig. 1A, Fig. S1A). Luciferase imaging was used to monitor growth of the primary tumor. Mice were then monitored weekly postoperatively to verify absence of gross disease and for recurrence (Fig. 1B). The two most common distant sites of pancreatic cancer recurrence in humans are the liver and peritoneal cavity, with isolated liver metastases occurring in approximately 25% of cases (*5*) (*10*). We found a similar frequency in our model, with 15% (n=6) exhibiting liver-only recurrence and 50% (n=20) with liver and peritoneal recurrence, both of which occurred early after surgery (mean time to recurrence 26 days) (Fig. 1C, Table 1). Another cohort (n=14; 35%) experienced latent recurrence with a mean time to recurrence of 554 days (Fig. 1C left, Table 1). Immunostaining for mCherry (Fig S1B) confirmed the origin of recurrent tumors as having arisen from the resected primary tumor. While not all of the latent cohort died over the course of the experiment (140 days), recurrent cancer was detected in 70%. Sites of recurrence in the latent recurring group included the liver, lung, perinephric fat, and uterine lining (Fig. S1C). Kaplan-Meier overall survival curves for the two murine cohorts exhibited striking similarity to survival curves in a cohort of 82 patients who had undergone surgery for PC and stratified into short term (median 0.8 years) and long term (median 6 years) survival groups (p<0.0001, Fig. 1C right) (*11*). There was a steep decline in survival early after resection that leveled out by approximately 60 days with 14 mice (35%) surviving beyond 100 days (Fig. 1D left panel). Median survival was 26 days (Fig. 1D left panel), which is equivalent to approximately 27 months in humans (*12*). Analysis of the patterns of overall survival amongst 41,552 stage I and II PC patients who underwent surgical resection (data from the National Cancer Database) showed similar characteristics with an early decline within the first 20 months that leveled out by approximately 50 months, and a median overall survival of 20 months (Fig. 1D right panel). Similar outcomes have been published from large single institutional databases (*5*) (*13*). We conclude that this model faithfully recapitulates the outcomes of human early stage patients undergoing resection for PC in both the location and frequency of distant relapse as well as overall survival.

**Figure 1.**
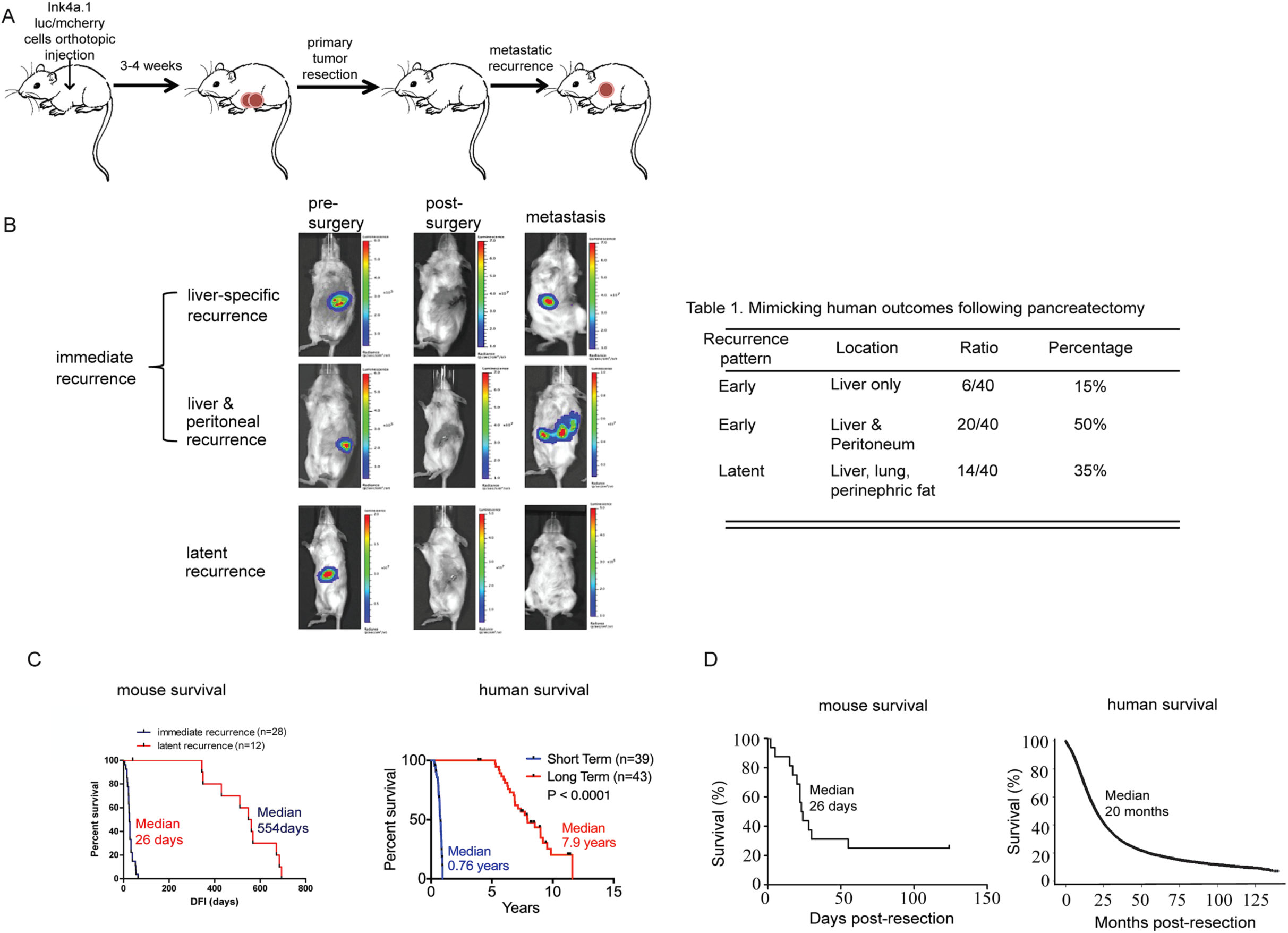
A novel model of early stage resectable pancreas cancer that mirrors human outcomes. **A)** Ink4a.1/luc/mcherry cells were injected into the tail of the mouse pancreas. Following 28-35 days post-cell injection, FVB mice underwent distal pancreatectomy with splenectomy. Mice would then recur as early as 5 days post-surgery, while another cohort would undergo latent recurrence. **B)** Mice were imaged using the IVIS spectrum following IP injection of luciferin (150 mg/kg) for cell detection prior to resection. Mice were imaged 3 days post-pancreatectomy and splenectomy. Mice had a disease-free interval (DFI) of 23 days (top), 17 days (middle), and 436 days (bottom) post-surgery (metastasis). Bar graphs indicate the range of radiance (photons/sec/cm2). Table 1: The ratio of early to latent recurrent and location site of recurrence in the resectable PC mouse model. **C)** Mouse and human survival segregating the early and latent recurrence groups. (Left) The median survival of mice from the short-term group was 26 days and the latent recurrence group was 554 days post-surgery, n=40. (Right) Survival of short- and long-term survivors of surgically resected pancreatic ductal adenocarcinoma patients in the MSKCC clinical cohort, n=82. **D)** Overall survival following primary tumor resection (left, n=40) and human (right, n= 49,555) Human data was acquired using the National Cancer Database for all resected PDAC patients with T1-3, N0-1 cancers.

The histology of resected primary tumors displayed a mixed morphology that contained a predominant mesenchymal phenotype with scattered glandular appearing cells (Fig. S2A*i*). E-cadherin staining of primary tumors exhibited a lack of staining in the mesenchymal cells and positive staining in the well-differentiated ones (Fig. S2A*ii*). Collagen deposition was observed at the stromal border along the tumor edge (Figure S2A*iii*). CD68 staining (Figure S2C*i*) confirmed exclusion of immune cells from the primary tumor, recapitulating the “cold” tumor microenvironment frequently seen in human PC. These features are consistent with previous classification of the Ink4a.1 cell line as a quasi-mesenchymal molecular subtype (*14*). Since the liver is the most common site of metastasis in PC, we focused our efforts on characterizing the recurrent tumors from the “liver-only” early cohort. Unlike the primary tumors, liver metastases displayed hallmarks consistent with the ‘classic’ molecular subtype of PC. This included more glandular tumor cells (Fig S2B*i*), with high expression of E-cadherin (Fig S2B*ii*) and robust collagen stroma (Fig S2B*iii*). Similar to the primary tumors, immune cells in liver metastases were mostly seen at the periphery (Fig. S2C*ii*). This apparent shift in features is consistent with the concept of mesenchymal-to-epithelial transition (MET) as playing a role in metastatic seeding (*15-18*). However, considerable heterogeneity was observed between metastatic liver lesions, even within the same mouse. Indeed, poorly differentiated metastatic tumors that retained the more mesenchymal features of the primary tumor were concomitantly observed along with those of a more glandular nature, with low to intermediate levels of E-cadherin, scant collagen deposition, and immune cell exclusion (Figure S2D). These data indicate that metastatic tumors arising from early recurrent mice exhibit phenotypic changes indicative of MET.

We hypothesized that the observed plasticity in morphology in the metastatic tumors is a consequence of epigenetic reprogramming, similar to that reported in human metastatic PCs (*19*). We examined a cell line derived from an early recurrent metastatic liver tumor for open chromatin regions in comparison to the parental cells using the assay for transposase accessible chromatin (ATAC-seq) (*20*). While genome-wide higher order chromatin accessibility patterns were largely comparable (see example of chromosome 1, Fig S3A), a total of 238 loci showed significant changes in chromatin accessibility. The primary tumor cell line, Ink4a.1, had 51 regions of open chromatin which were not observed in the metastatic line, Met38, whereas the latter had gain of open chromatin in 187 regions (Fig. S4). Examples of genes with more open (Elf3) and closed (Oxr1) regions in metastatic cells are shown in Fig. S3B-C. Taken together, our data suggest that the metastatic behavior of this model system exhibits pathological, histological, and chromatin-level features similar to human PC metastasis. This further supports the use of this model system for investigating metastatic tumor cell dormancy.

### Latent recurrent mice harbor disseminated tumor cells originating from the primary tumor

As our model displays a cohort of mice that with a very long latency period, we sought to determine if we could use these mice to isolate disseminated tumors cells from the liver for studying dormancy. Given the rarity of this cell population in the liver, it was important to first validate that we could isolate these cells and verify they originated from the primary tumor. We used fluorescence activated cell sorting (FACS) to sort mCherry^+^ cells (Fig. S5A) from the liver of a mouse with a DFI>200 days (whole body MRI confirmed absence of radiographically detectable disease, Fig. S5B). We isolated genomic DNA from the sorted mCherry^+^ cells as well as several controls and amplified with primers specific for luciferase, Kras^WT^, and Kras^G12D^. The luciferase gene was present in the sorted mCherry^+^ cells (DTCs) from this mouse as well as in the unsorted parental Ink4a.1 cells, and sorted mCherry^+^ cells from mouse that received an injection of 1×10^6^ cells intrasplenically. We used liver DNA taken from an uninjected mouse as a negative control (Fig S5C). We also detected Kras^G12D^ specific DNA in the sorted mCherry^+^ cells from a mouse with DFI>200 days, as well as the Ink4a.1 cell line and Kras^G12D/+^ tail DNA, but not in the uninjected liver, KRAS^WT^ tail DNA, or no DNA sample (Fig. S5D).

As an orthogonal method of validating the mCherry^+^ sorted cells were from the primary tumor we turned to transcriptomic profiling using single cell RNA sequencing because it can distinguish tumor and different non-malignant cell types at single cell resolution and also provide insights into the transcriptomic makeup of DTCs and heterogeneity thereof – providing a distinct advantage. Jointly analyzing single cell expression data using principal component analysis from mCherry^+^ flow sorted cells from the primary tumor (#324, n= 116) and DTCs from the liver (n=109 and 110 from two individual mice #239, #241), and also normal liver tissue from the Tabula Muris (*21*) (see Supplemental Methods) we observed that the flow-sorted cells on average were highly similar to the primary tumor cells and very different from the normal liver cells (Fig. S5E). Taken together, these observations support that a vast majority of the isolated mCherry^+^ cells harvested using flow cytometry from latent recurring mice are indeed DTCs that originated from the primary tumor.

It has been speculated that one reason latent recurrent patients survive longer is that they harbor fewer DTCs then their short surviving counterparts. There is evidence that the burden of DTCs correlates with survival in certain cancers (*22*) (*23*). We examined this by quantifying the DTC burden in the liver at the time of pancreatectomy and comparing this to the DTC burden in dormant mice. We detected between 0.01-0.04% mCherry^+^ cells at the time of primary tumor resection (Fig S6A-F). Interestingly, the frequency of mCherry^+^ cells in the livers of dormant mice (DFI>100 days post resection) was determined to be between 0.01-0.1%, a number that was significantly more than at the time of surgery (Fig. S7A). Thus, we conclude that the latent recurrent phenotype cannot be explained simply by a lower burden of minimal residual disease. The higher number of DTCs present in dormant mouse liver may be due to reactivation of single cells into micrometastases that activate the immunogenic dormancy response (*24-26*).

Patients that suffer metastatic recurrence after surgery from pancreatic cancer demonstrate several common sites of recurrence (liver, abdominal lymph nodes, peritoneum, lung); it is unclear to what extent they harbor DTCs in other organs. To illuminate this, we calculated the frequency of DTCs in the bone marrow, brain, heart, intestine, kidney, liver, lung, and stomach from three dormant mice (Fig. S7B). Rates were exceedingly variable (0-0.6%, n=3) but every organ presented with DTCs depending on the mouse. The greatest organ dissemination frequency was not the same for all: #239 and #241 had more in the stomach and liver, while #249 had more in the intestine and heart. However, all mice harbored detectable DTCs in the bone marrow, heart, and liver demonstrating that DTCs were ubiquitous throughout the mouse despite the fact that latent recurrences were not ubiquitous but rather occurred mostly in the lung and liver. This suggests that the microenvironment of the DTC plays an important role in reactivation from dormancy.

### Transcriptomic profiling reveals the molecular pathways that govern pancreatic cancer dormancy and identifies a dormancy gene signature

Our first approach to investigate the biology of pancreatic cancer dormancy was to examine the transcriptomes of cells using both standard (ultralow) and single cell (10X Genomics) RNA-seq methods. We compared the transcriptomes from several cell populations (Fig. 2A), including the primary tumor, DTCs isolated from the livers of latent recurrent mice (dormant DTCs), and proliferating cells that were cultured in the presence of 10% serum from DTCs harvested from the livers of latent mice (hereafter called ‘reactivated clones’). These latter cells remained quiescent for months (average 190 days, n=4) before spontaneously resuming proliferation at which time we harvested them for RNA after no more than 3 cell passages. This resumption of proliferation is one indication that these cells were not irreversibly senescent. We had difficulties in obtaining RNA from latent recurrent tumors in mice due to the lack of predictability and rarity of this event, as well as the aggressiveness of pancreatic cancer recurrence.

**Figure 2.**
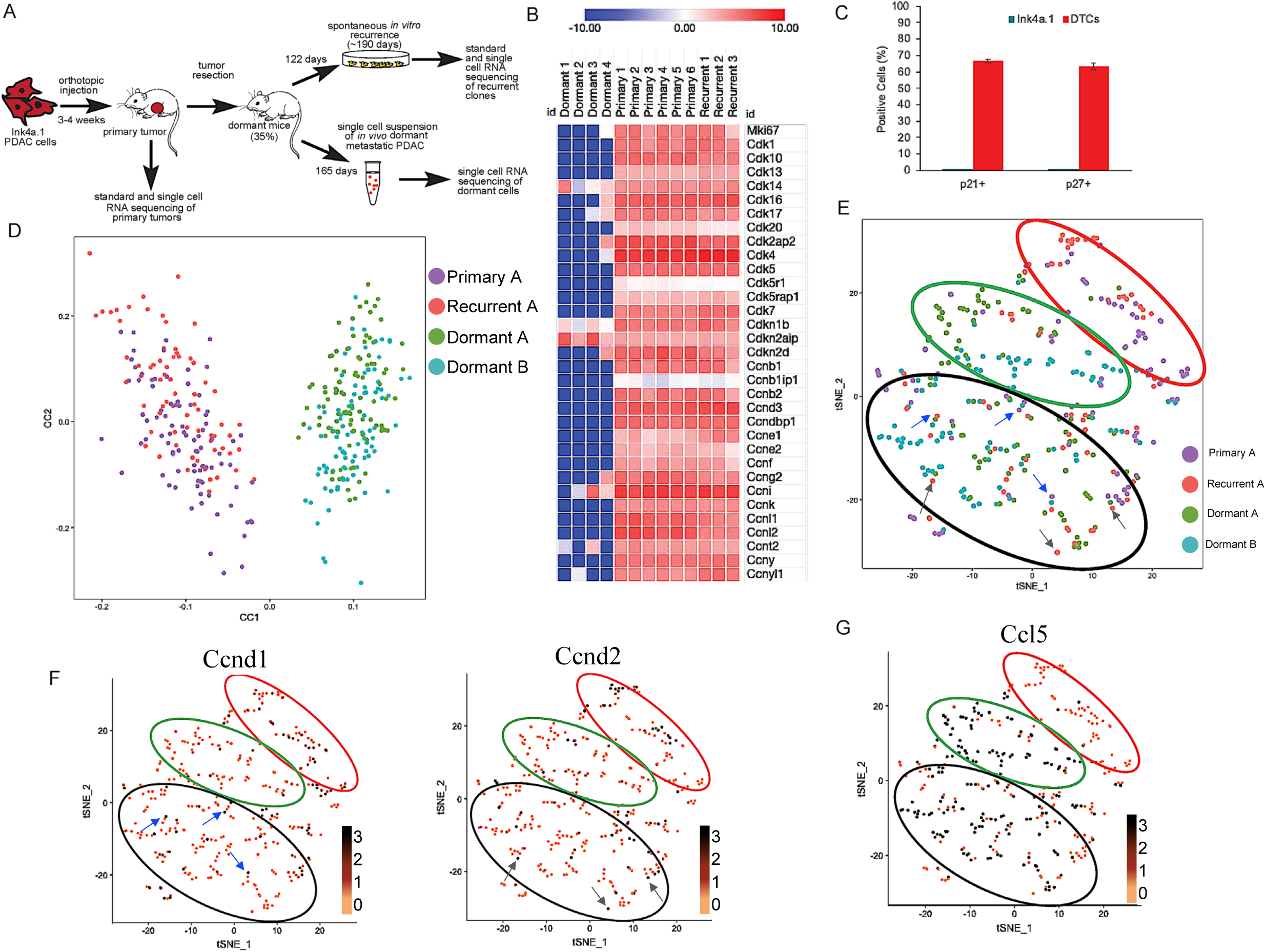
Single cell transcriptomic profiling reveals the plasticity and heterogeneity of pancreatic cancer dormancy. **A)** Schematic representation of sample origins used for standard and single cell RNA sequencing. **B)** Expression of markers of cell cycle analyzed from ultralow RNA-seq analysis of individual dormant DTCs (dormant 1-4) compared with primary tumors (Primary 1-6) and *in vitro* recurrent cells (Recurrent 1-3). Transcript expression was calculated using the log_2_ values of the significant non-zero values of TPMs. Heat maps were created using Morpheus from the Broad Institute at https://software.broadinstitute.org/morpheus/ Flow cytometry analysis of cells positive for p21 and p27 expression. The parental primary tumor cell line Ink4a.1 was used as a control. DTCs from the livers of dormant mice were analyzed for expression of CD44, CD133, mCherry, p21, and p27 expression. The parental primary tumor cell line Ink4a.1 was used as a control. Results shown are for CD44^+^/CD133^+^/mCherry^+^/p21^+^ or CD44^+^/CD133^+^/mCherry^+^/p27^+^ cells. **D)** CCA plot of expression data from #324 (primary tumor A), #226 (recurrent clone A), and the dormant DTCs from mice #239 and #241 (Dormant A, Dormant B). **E)** tSNE plot showing the distribution of clusters from expression profiles of primary tumor (purple), recurrent tumor (red), and dormant DTCs from two separate mice (green, blue). Cluster I: red circle, Cluster II: green circle, Cluster III: black circle. Blue and gray arrows represent cells with high expression of Cyclin D1 and D2, respectively. **F)** Expression of proliferation markers Cyclin D1 (left) and Cyclin D2 (right) in tSNE plots displaying high expression in Clusters I and III. log_2_TPM expression shown as a gradient. Blue and gray arrows as in **E. G)** Expression of possible dormant DTC marker Ccl5 in the tSNE plots as in **(E)** displaying high expression in Clusters II and III.

We first sought to determine if the DTCs from latent recurrent mice exhibited features associated with quiescence. We first used gene expression data from the ultralow input protocol. We sequenced three individual cells (Dormant 1-3) as well as a pool of 100 cells (Dormant 4). To determine quiescence, we examined markers of proliferation including Ki67 and a large panel of cyclins and cyclin dependent kinases. We found that expression of proliferation genes by dormant samples was decreased in comparison to primary tumor and recurrent samples (Fig. 2B). Since cdk inhibitors p27 (*Cdkn1b*) and Carf (*Cdkn2aip*), a regulator of p21, were shown to be expressed in dormant DTCs (Fig. 2B, (*27*) (*28*)), we verified that dormant DTCs obtained from livers of dormant mice were upregulating p21 and p27 (Fig. 2C). Thus, we concluded that these DTCs are of a dormant phenotype.

We sought to understand the biological pathways that may be important during dormancy by analyzing genes and pathways significantly differentially expressed between dormant DTCs, primary tumors, and reactivated clones using the both the 10X Genomics (Table S1) and the standard ultralow (Table S2) data sets. We constructed a dormancy signature comprised of significantly differentially upregulated genes (fold change >2; FDR adjusted p-value < 10^−4^) between the dormant DTCs (#239, #241) versus the primary tumor (#324) and reactivated clones (#226). An analysis of upregulated genes reveals possible immunoregulators, such as *Ccl5, Btnl9*, and *Xcl1*, multiple C-type lectin family members (*Clec4f, Clec1b, Clec14a*), genes involved in the metabolism or transport of lipids (*Apoe, Apoa1, Apoa2, Apoc1, Fabp1*), and genes that function in angiogenesis (*Kdr, Adgrl4, Egfl7*). Many genes associated with the integrin signaling pathway, such as *Col5a2, Fn1, Col8a1, Actg2*, were downregulated (Table 2).

**Table 2.**
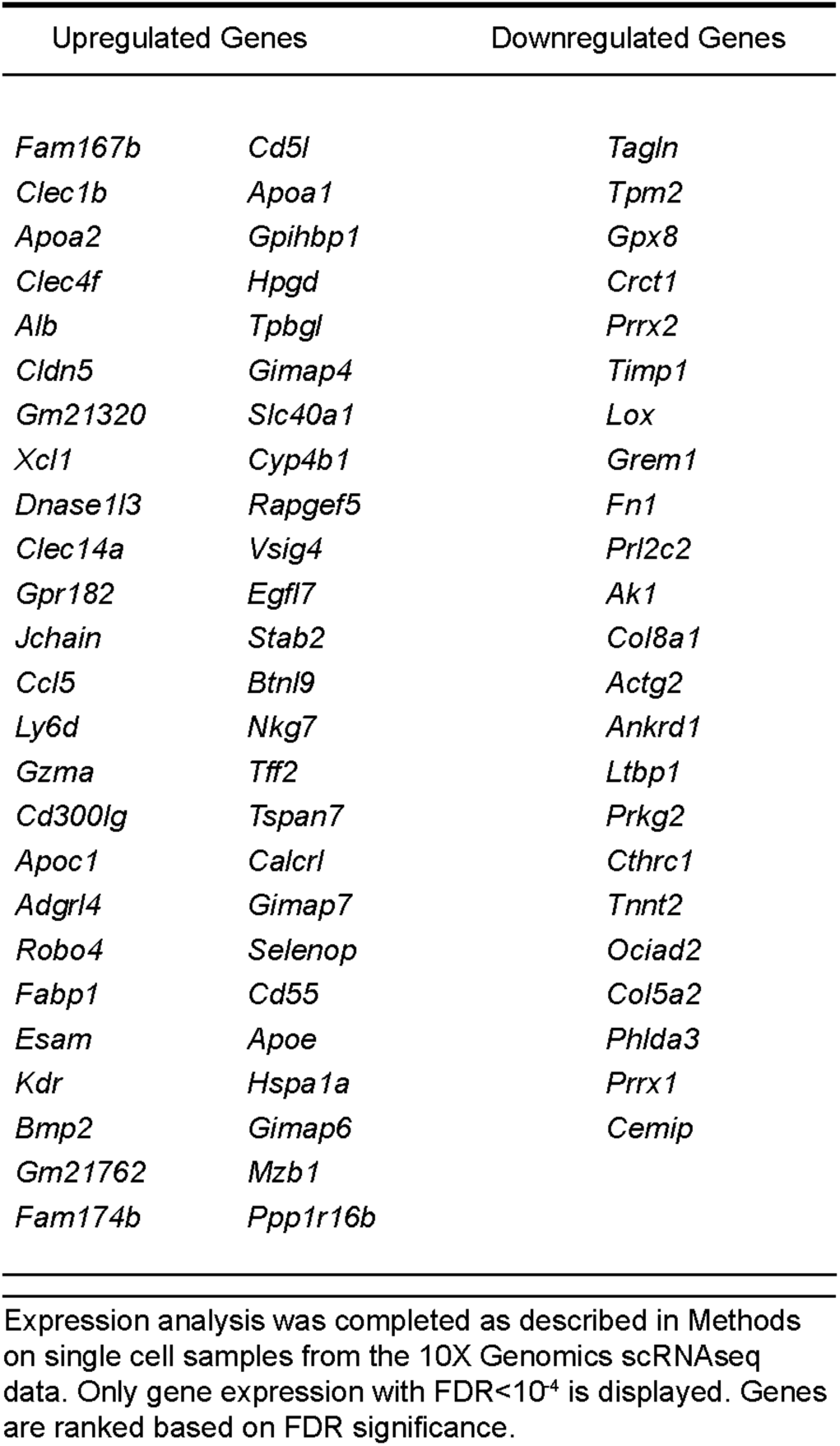
Dormancy signature in metastatic pancreatic cancer. The top differentially expressed genes that were up- or down-regulated were used to create a gene signature from the 10X genomics data. Expression analysis was completed as described in Methods. Only gene expression with FDR<10^−4^ is displayed. Genes are ranked based on FDR significance.

The immunomodulators expressed in the DTCs include checkpoint inhibitors (*PDL1, CD38, Btnl9*), cytokines (*IL1, IL6*), chemokines (*Ccl28, Ccl5, Cxcr4*), and receptors (*GMCSFR, IL6R*), suggesting that dormant DTCs are capable of immunosuppression in the tumor microenvironment (*29-31*). Calcium signaling, through upregulation of *Mcu* and *Ncx*, may be used for energy maintenance as well as ER stress and ROS production control (*32*), while increased linoleic metabolism (*Cyp2c, Alox15, Pla2g4a, Cyp2j*) may provide linoleic acid metabolites that function in cell signaling (*33*) (*34*) (*34*). Retinoic acid has been shown to be important in maintenance of dormancy in hematopoietic stem cells (*31*), breast cancer, and squamous cell carcinoma (*28*). Many neuroactive receptor genes were overexpressed, including the Neuropeptide Y receptor that functions to modulate circadian rhythm (*35*) (*36*).

At the pathway level, our analysis revealed cellular processes involved in the deactivation of the immune system (Figs. S8-S9), neuroactive ligand-receptor interaction (Fig. S10), metabolism of retinoids and linoleic acid (Fig. S11), olfactory transduction, Jak-STAT signaling, calcium signaling, PPAR signaling, ABC transporters, drug metabolism, cell adhesion molecules, and bone morphogenic proteins (BMPs, Fig. S8 arrows). Accordingly, pathways most affected by gene downregulation in both data sets revealed a cell program of decreased cell proliferation, protein biosynthesis/expression, and cellular energy metabolic pathways, such as the TCA cycle, oxidative phosphorylation, and the pentose phosphate pathway (Table S4). Each single cell RNA-seq protocol gave a variety of pathways, possibly due to technical differences in the protocols. However, there were pathways that were homologous between the sets of data (Table 3). Taken together, DTCs exhibit a dormancy gene and pathway signature marked by immune regulation, cell adhesion, cell cycle, calcium signaling, and metabolic processes.

**Table 3.** Homologous Pathways in Dormant Cells. Ultralow and 10X genomics data was normalized using edgeR and pathway enrichment completed using GAGE (see Methods for more detail). Pathways present in both data sets are listed here, with q<0.1.

| Pathway Name | q value (UL) | q value (10X) |
| --- | --- | --- |
| <u>Upregulated Pathways</u> |  |  |
| olfactory transduction | 4.80E-124 | 2.29E-04 |
| neuroactive ligand-receptor interaction | 1.41E-31 | 1.50E-03 |
| cytokine-cytokine receptor interaction | 4.13E-04 | 6.79E-08 |
| hematopoietic cell lineage | 4.13E-04 | 1.21E-07 |
| primary immunodeficiency | 5.38E-03 | 2.09E-07 |
| retinol metabolism | 1.08E-02 | 8.55E-03 |
| cell adhesion molecules | 1.75E-02 | 1.52E-06 |
| intestinal immune network for IgA production | 1.79E-02 | 4.52E-08 |
| graft-vs-host disease | 1.87E-02 | 2.21E-12 |
| calcium signaling pathway | 1.02E-02 | 5.56E-02 |
| linoleic acid metabolism | 4.18E-02 | 5.22E-02 |
| African trypanosomiasis | 7.25E-02 | 6.89E-03 |
| <u>Downregulated Pathway</u> |  |  |
| cell cycle | 6.60E-08 | 4.34E-02 |

q<0.1 for all pathways. UL=ultralow input RNAseq dataset, 10X=10X Genomics input RNAseq dataset

### Single cell transcriptomic profiling reveals the plasticity and heterogeneity of pancreatic cancer dormancy

An active question in the metastasis/dormancy field is whether dormant tumor cells are a subpopulation of primary tumor cells that have the capacity to become dormant and are selected before dissemination, or is the dormant phenotype acquired after dissemination (*37*). The latter model would suggest that dormancy and reactivation is a plastic process of transdifferentiation. To shed light on this using the 10X Genomics dataset we performed a canonical correspondence analysis (CCA) where mouse #324 (primary tumor A) and mouse #226 (reactivated *in vitro* clone) cluster together, and away from the dormant DTCs from mice #239 and #241 (Dormant A and Dormant B) which also clustered together (Fig. 2D). This analysis supports a model of plasticity in dormancy where the primary tumor cells differentiate into dormant tumor cells and then revert back upon reactivation. We next focused on the question of cell to cell heterogeneity among dormant tumor cells using the 10X Genomics data. To examine the degree of heterogeneity, we used t-distributed stochastic neighbor embedding (tSNE) as implemented in the Seurat pipeline (*38*) to analyze cell population-level heterogeneity and to identify sub-populations in the primary tumor (#324), reactivated clone (#226) and dormant DTCs (#239, #241). Again, cells from the primary tumor and that from the reactivated clones more closely clustered together and away from the two dormant samples (Fig. 2E) Interestingly none of the four samples completely segregated from one another, suggesting a fair degree of intragroup heterogeneity and group overlap in the transcriptomes of different cell populations. We observed three major subpopulations of cells in the tSNE plot – Cluster I (red circle): dominated by cells from primary tumor and reactivated clones, Cluster II (green circle): dominated by dormant DTCs, and Cluster III (black circle): mixed. This mixed cluster suggests that perhaps there is a transient state between the proliferative and dormant states. When we overlaid expression of cell cycle marker genes such as *Ccnd1*and *Ccnd2*, Cluster I showed higher expression of these genes while that of Cluster II showed lower expression (Fig. 2F). Interestingly, those cells that showed higher cell cycle gene expression in Cluster III mapped to cells from the primary tumor or to reactivated clones (compare blue and grey arrows in Fig 2E to corresponding blue and black arrows in Fig. 2F), again showing the relationship between proliferative activity and cells of the primary tumor and reactivated clones.

Recently the chemoattractant cytokine CCL5 has been implicated in hepatocellular carcinoma dormancy through an immune escape mechanism by recruitment of regulatory T cells (T regs) (*39*). CCL5 is a gene we found upregulated in the dormancy gene signature. We found high CCL5 expression in Cluster II, low expression in cluster I and high expression in Cluster III mapping to the cells of the dormant DTCs samples (Fig. 2G). This indicates that CCL5 is a novel dormancy marker in this murine model.

### The murine dormancy gene signature has relevance to human pancreatic cancer patients

To determine if the murine dormancy gene signature has relevance in human pancreatic cancer, we performed a parallel analysis on DTCs from two human pancreatic cancer patients. Using an IRB approved clinical protocol, samples of primary tumor and normal appearing liver were collected during surgery, made into cell suspensions and both samples were sorted by FACS using an antibody to CA19-9. We then performed single cell RNA sequencing (10X Genomics) on the CA19-9^+^ populations from the primary tumor and liver (Fig. 3A; See Method for details). The first patient had 147 and 262 cells profiled from the primary tumor and liver, respectively, while the second patient had 73 and 211 cells profiled from the primary tumor and liver, respectively. FACS-enriched cells had tumor as well as non-tumor cell populations. The cell population consisting of primary and disseminated tumor cells was identified by gene signatures using a published approach, and the results were visualized on a t-SNE plot constructed from the principle components (Figure 3B). The first patient had 47 and 22 potential tumor cells from the primary tumor and liver, while the second patient had 44 and 58 potential tumor cells from the primary tumor and liver, respectively, suggesting that the CA19-9^+^ cell population from the liver had reasonable representation of DTCs (8.4% in patient P3552 and 27% in patient P7180).

**Figure 3.**
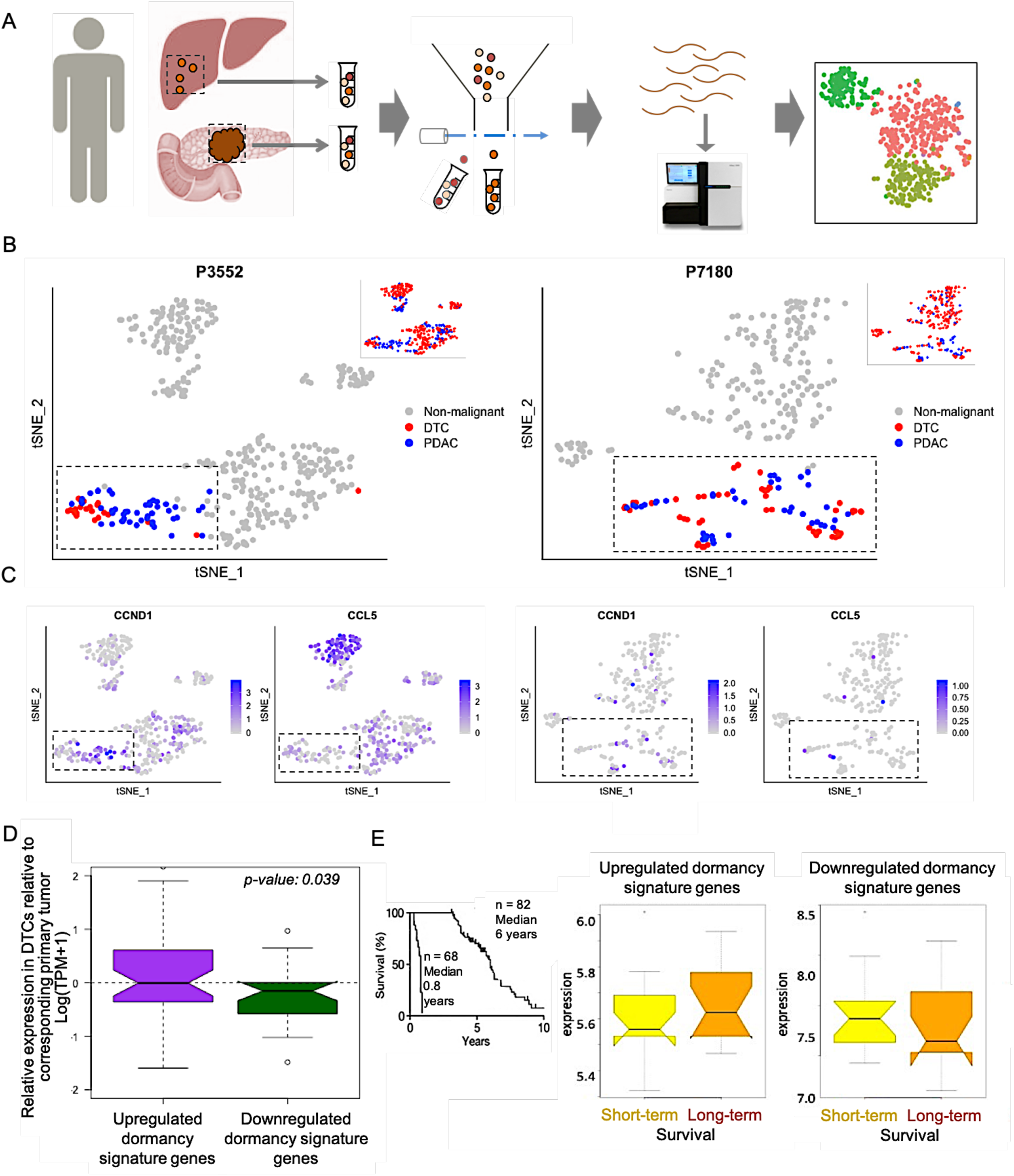
Clinical relevance of the dormancy signature derived from the transgenic mouse model in human pancreatic adenocarcinoma. **A**) Schematic outline of single cell RNA sequencing of CA19-9 positive cells from tumor and liver from pancreatic cancer patients. **B**) tSNE plots show projection of primary tumor (blue) and liver derived (red) CA19-9 positive cells from 2 patients (P3552 and P7180). Insets show all isolated cells, and the main panels show the potential tumor cells after excluding non-tumor cells based on gene expression signature-guided annotation. **C**). tSNE plots showing single cell expression of *CCND1* and *CCL5* in primary tumor and liver derived CA19-9 positive cells from the two patients. Only the potential tumor cells are highlighted. **D**) Boxplot showing differences in log_2_(TPM) expression between DTCs and primary tumor in human for the dormancy signature genes up- and downregulated in mouse DTCs. p-value was calculated using one-tailed Wilcox rank sum test. **E**) (Left) Kaplan-Meier survival curves for the early and latent recurrent human patient cohorts used to derive gene expression data on right. (Right) Boxplot showing differences in log_2_(TPM) expression of the dormancy signature genes upregulated and downregulated in mouse DTCs in the patients who have early and late relapse.

We then examined the expression of cell cycle marker genes such as *CCND1* and found high expression in cells from the primary tumor, but lower expression in the DTCs and negligible expression in normal liver cells, furthering the support that the DTC population is predominantly quiescent (Fig. 3C). Consistent with our observations in mouse DTCs, we also found *CCL5* was expressed at a significantly higher level in the human DTC population as compared with the primary tumor or liver populations, validating CCL5 as a marker for dormant DTCs in both murine and human systems (Fig. 3C).

We next sought to determine if the dormancy gene signature we identified in the mouse model could discriminate the DTC population in human pancreatic cancer patients. We computed the differences in log_2_(TPM) expression between DTCs and primary tumor for the dormancy signature genes inferred from the mouse model. We found that the genes that are up- and downregulated in mouse DTCs show significantly positive and negative differences, respectively, in expression between patient DTCs and primary tumor – indicating that upregulated dormancy signature genes are also highly expressed in human DTCs, while downregulated dormancy signatures show the opposite pattern (p<0.039, Wilcox Rank Sum test) (Fig. 3D). Therefore, not only are select biomarkers, such as cyclins and Ccl5, relevant for discriminating human DTCs, but the entire 73-gene dormancy gene signature is also suitable to determining DTCs.

We then analyzed the murine dormancy signature in human PC primary tumor specimens from patients with short and long-term survival to determine if the signature could predict short-term versus long-term survival after surgery. We compared specimens from 15 short-term survival patients (median survival 0.8 years) and 15 long-term survival patients (median survival not yet reached) (Fig. 3E, left) (*11*) and assessed whether the dormancy signature genes showed a differential pattern of expression between the two groups (Fig. 3E, right). Interestingly, among the dormancy signature genes, those upregulated were expressed more highly in long-term survivors as compared to the short-term survivors. Likewise, the genes that were downregulated in the signature were lowest in expression in the long-term survivors versus the short-term survivors. However, these data were not statistically significant (p=0.18 and 0.17, respectively). We also evaluated a separate cohort of primary tumors (n= 39 long-term, n=43 short-term survivors), and once again observed that genes upregulated in the dormancy signature were more highly expressed in the long-term survivors, and genes downregulated in the dormancy signature were less expressed. The lack of statistical significance could be a result of multiple circumstances. First, the RNA specimens from the human primary tumors were extracted from tumor tissue and are thus a heterogeneous population of non-tumor and tumor cells (as suggested by Fig 2E, Cluster III), with a low proportion of dormant cells. This could cause a dilution of the dormancy signature in transcriptomic data. Secondly, the microarray expression data did not contain the entire 71 gene signature due to lack of gene inclusion on the chip and/or degraded RNA from the FFPE samples. Alternatively, the dormant phenotype may begin in some cells in the primary tumor but is fully completed upon dissemination and colonization.

### Pancreatic Cancer Dormancy is characterized by global chromatin remodeling

Given the transcriptomic changes we observed in isolated DTCs relative to the primary tumor and/or *in vitro* reactivated clones, we applied the ATAC-seq assay to compare chromatin accessibility between 4 *in vivo* samples: two liver-derived dormant DTC populations each harvested 451 days after surgery were compared to one recurrent peritoneal tumor found 471 days after surgery processed in duplicate. For the initial analysis, the aligned reads were down sampled to the same number for all 4 data sets (52.8 million in this case), and these normalized files were used for peak calling. Merged Peak Regions are the union of the 4 peak sets, and thus are defined as all genomic regions where at least one of the 4 samples showed a called peak. The analysis identified 17,737 such regions. A comparison of the distribution of tag depth (aligned reads) at each of these Merged Regions among the 4 samples showed an overall lower peak signal in the dormant samples relative to recurrent tumors, suggesting significantly fewer peaks and/or a broad decrease in signal intensity (Fig. 4A). To identify regions that displayed changes that significantly deviate from this global effect, we applied the DESeq2 algorithm. This method first normalized the 4 data sets such that the overall tag distribution in the peak regions is the same. Using a cutoff of FDR <0.1, DESeq2 identified 1692 sites that were differentially accessible; 1310 that were more accessible in the DTCs than the recurrent tumors and 382 that were less accessible. A heatmap representing 296 most differentially accessible sites is shown in Fig. 4B. The genes within these sites represented in this map are listed in Table S4.

**Figure 4.**
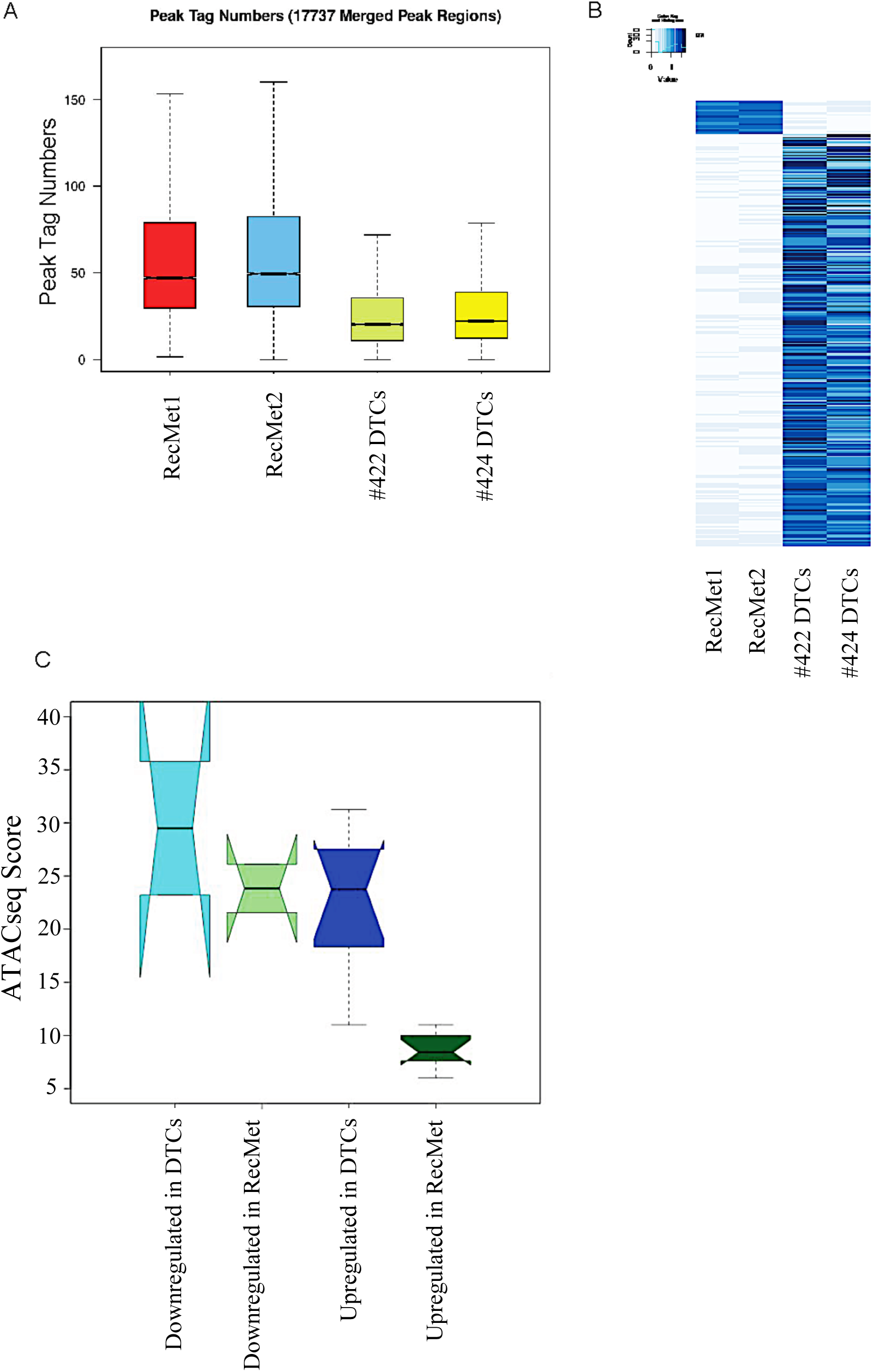
Pancreatic Cancer Dormancy is characterized by global chromatin remodeling. **A)** ATAC-seq boxplot analysis showing the peak tag numbers for a recurrent tumor processed in duplicate versus two dormant DTC samples from two separate mice. **B)** ATAC-seq heatmap of genes in open (dark blue) and closed (light blue) regions for a recurrent tumor processed in duplicate versus two dormant DTC samples from two separate mice. The data was analyzed as described in Methods. **C)** Correlation of the genes up and down regulated in the murine dormancy gene signature with ATAC-seq score.

We further investigated whether the dormancy signature genes significantly up and down regulated in dormant DTCs displayed changes in chromatin accessibility. The gene sets showed overall increased chromatin accessibility, consistent with genome-wide patterns. A minor subset of dormancy genes was associated with significant peaks in the ATAC-seq assay. While those down regulated showed no significant difference in chromatin accessibility, up-regulated genes showed a significant increase (FDR adjusted p-value < 0.05) in chromatin accessibility in DTCs, consistent with their increased expression (Fig. 4C, 4D). Taken together, we conclude that the dormant cancer state is characterized by widespread changes in chromatin accessibility.

### Dormant pancreatic cancer cells adopt a stem cell phenotype that is lost upon reactivation

It has been hypothesized that the dormant tumor cell population is enriched for cancer stem cells (CSCs) but it is not known if this is because these cells are selected for during metastatic progression or if these properties are acquired after dissemination (*37*). Using flow cytometry for cell surface markers and the aldefluor assay for aldehyde dehydrogenase activity (ALDH), we quantified the expression of several markers of pancreatic cancer CSCs. We compared cell populations consisting of parental Ink4a.1 cells (primary), flow sorted dormant DTCs from the liver (dormant), a cell line made from a liver metastasis from an early recurrent mouse (early recurrent), and reactivated clones. The primary tumor cell EpCAM expression was extremely low (<1%) with moderate c-Met (∼25%) expression, consistent with the QM molecular subtype of these cells (Fig. 5A). Expression of CSC markers in the primary tumor cell line was more heterogeneous: 100% CD44^+^, ∼30% Aldh1^+^, 5% CD133^+^ and 20% CD24^+^. DTCs from the livers of dormant mice expressed even higher levels of CSC markers (100% CD44^+^, 100% Aldh1^+^, and 80-100% CD24^+^, 90-100% CD133^+^, and 100% c-Met^+^. Cells from the early recurrent liver metastasis and reactivated clones were low in expression of many of the CSC markers: Aldh1, CD44, CD133, and c-Met. Interestingly these reactivated clones expressed high levels of EpCAM, consistent with the MET observations we made in liver metastases of early recurrent mice (Fig. S2B). CD24 expression was high (>80%) in all liver metastatic groups: dormant, early recurrent and reactivated clones, demonstrating the role of P-selectin in metastasis (*40*). We then examined the single cell transcriptomic data for further evidence of a stem cell phenotype. BMPs have been shown to regulate quiescence and long-term activity of neural stem cells (*41*) and dormancy of prostate cancer cells (*42*). We found high expression of several BMP family members (BMP2,4-7) in the dormant population of cells (Fig. S8, arrows). When we analyzed BMP2 expression at the single cell level by tSNE plot, we observed highest BMP2 expression in Cluster II (green circle, dormant population) and Cluster III (black circle, mixed population), indicating dormant DTC expression. The lowest expression was observed in Cluster I (red circle) which was dominated by proliferating cells from the primary tumor and reactivated clones (Fig. 5B).

**Figure 5.**
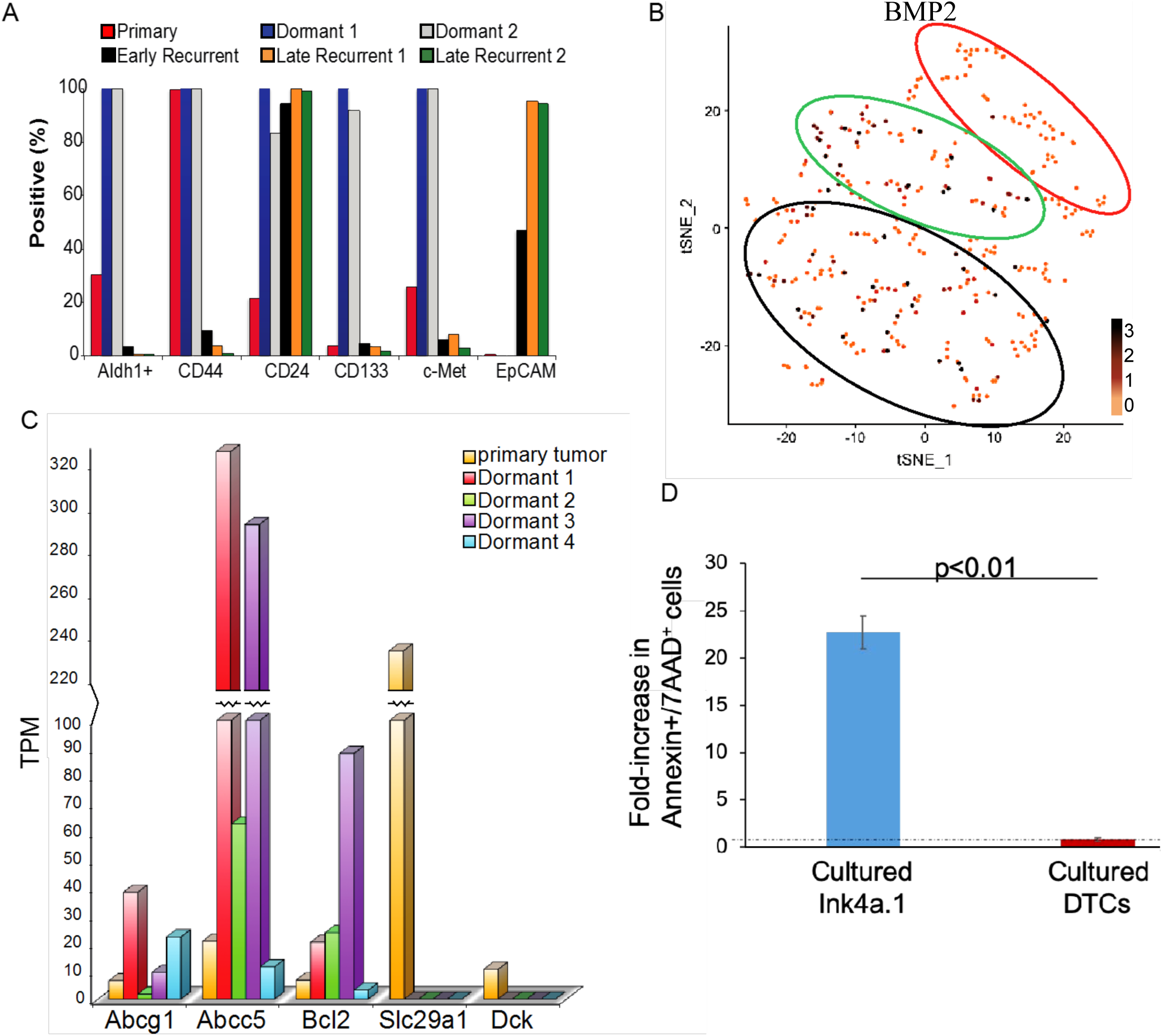
Pancreatic Cancer DTCs adopt a stem cell-like phenotype. **A)** Primary pancreatic cancer cell line Ink4a.1, dormant mCherry+ cells harvested from the liver of two separate mice, early recurrent cell line, Met38, and latent recurrent cell lines were assessed by flow cytometry for their expression of CSC markers Aldh, CD44, CD24, CD133, c-Met, and EpCAM. **B)** Expression of dormant DTC marker *Bmp2* in the tSNE plots as in Fig. 2F displaying high expression in Clusters II and III. log_2_TPM expression shown as a gradient **C)** Expression of known chemoresistance markers in dormant cells was determined using the ultralow single cell RNA-seq protocol comparing expression of a primary tumor with single cell samples Dormant 1-4. **D)** Sorted metastatic pancreatic cancer DTCs and parental Ink4a.1 primary tumor cells were treated with 100 nM gemcitabine for 48 hrs *in vitro* and cell death assessed using Annexin V and 7-AAD staining.

As chemoresistance is a putative property of cancer stem cells (*43*), we tested the sensitivity to chemotherapy in the dormant DTCs as a functional assay to further substantiate their stem cell phenotype. Since these cells are dormant, we could not use other functional assays that examine tumor initiating capacity. We tested their sensitivity to gemcitabine *in vitro* using an assay for apoptosis as these cells are not cycling. Parental Ink4a cells and flow-sorted dormant tumor cells were exposed to 100 nM gemcitabine for 48 hours and then assayed for apoptosis using Annexin V staining. Gemcitabine induced a greater than 10-fold increase in apoptosis in the Ink4a cells as compared to the vehicle control whereas the dormant cells show little evidence of apoptosis over the vehicle, indicating that the dormant DTCs are gemcitabine-resistant (Fig. 5C). We then examined gene expression levels in our dormant DTCs for several genes that are associated with drug resistance including cell membrane transporters (*Abcg1, Abcc5*), cell survival (*Bcl2*), as well as gemcitabine import (*Slc29a1*, aka *hENT1*) and gemcitabine activation (*Dck*, deoxycytidine kinase). Using the data from the ultralow input RNA-seq we compared gene expression levels in the primary tumor cells to that of dormant cells (Dormant 1-4). We found an increase in expression of *Abcg1, Abcc5*, and *Bcl2* in the dormant samples and a decrease in expression in *Slc29a1* and *Dck*. This suggests that the mechanism of resistance is multi-factorial (Fig. 5D). Taken together these results support a conclusion that dormant pancreatic cancer cells undergo a differentiation change and adopt stem cell-like properties that are lost upon reactivation. Furthermore, these results are consistent with the plasticity we observed in the single cell transcriptomic data (Fig. 4B) and support a model in which these properties are acquired after dissemination rather than selected in the primary tumor.

### Mechanisms of Pancreatic Cancer Dormancy are cell autonomous and non-autonomous

Recently several transcription factors have been identified to regulate dormancy including the circadian rhythm gene *Bhlhe41* (Dec2) and *Nr2f1* (*2*) (*28*). We found in our dormant samples that Dec2 was significantly upregulated as compared to the primary tumor and then downregulated in the recurrent clones (Fig. 6A). We did not find an upregulation of *Nr2f1* in our dormant samples. As our transcriptomic analysis of the dormant DTCs identified the circadian rhythm gene Dec2 as upregulated in our dormant model, we sought to validate this as well as examine its functional significance. We validated our RNA-seq data by examining both Ki67 and Dec2 protein expression by flow cytometry and found cells isolated from the livers of latent recurrent “dormant” mice were Dec2^hi^ Ki67^lo^, whereas the cells from the primary Ink4a.1 cell line were the opposite (Fig. 6B). To determine if Dec2 was functionally relevant in dormancy we overexpressed Dec2 in the Ink4a.1 cells and performed co-immunofluorescence for Dec2 and Ki67. We found that in cells in which Dec2 expression was high, Ki67 expression was low, indicating Dec2 induced quiescence in these tumor cells (Fig. 6C).

**Figure 6.**
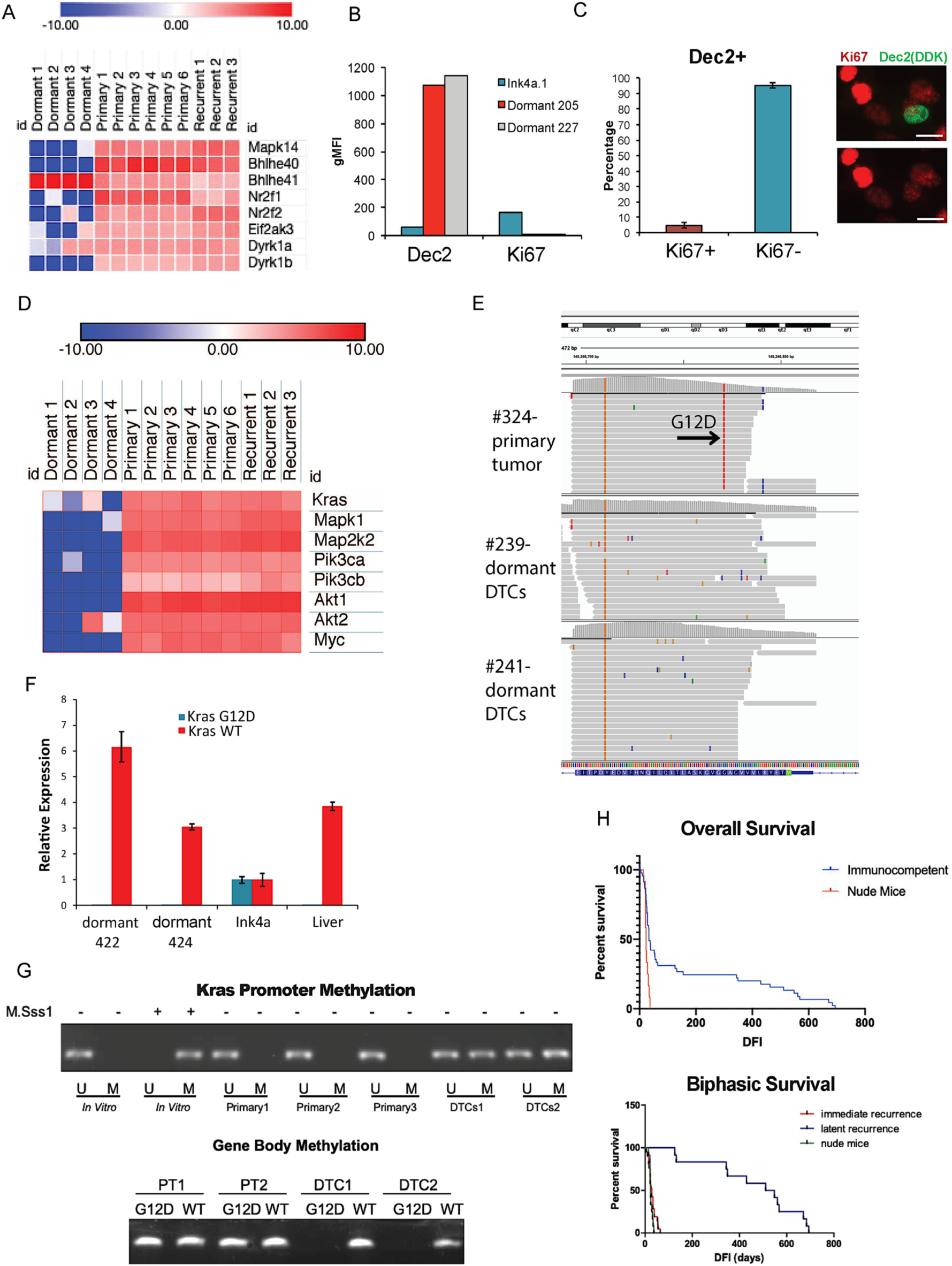
Mechanisms of Pancreatic Cancer Dormancy are cell autonomous and non-autonomous. **A)** Expression of dormancy markers as in Fig. 2B. **B)** Flow cytometry analysis of Dec2 and Ki67 expression in primary cell line Ink4a versus DTCs from 2 dormant mouse livers. The gMFI was calculated using FlowJo. **C)** (Left) Ki67 expression in Ink4a.1 cells expressing a Dec2-DDK/myc plasmid. (Right) Immunofluorescence staining of Ki67 and DDK(Dec2) (top) showing a lack of expression of Ki67 when Dec2 is expressed (bottom). Scale bar: 15 *μ*m.E. Expression of cell proliferation and apoptosis genes using the same methods as in B. **D)** Expression of Kras and proliferative markers as in Fig. 2B. **E)** IGV plot of the Kras locus focusing on the sequence surrounding the G12D mutation. Notice lack of G12D mutation (red vertical line) in the dormant RNA transcripts. **F)** Expression analysis of mutant Kras vs WT Kras completed using qPCR in dormant DTCs (DFI>400 days), Ink4a.1 parental cells (positive control), and liver cells. **G)** Methylation-specific PCR using primers located in the CpG island of the Kras promoter (top, U: unmethylated, M: methylated primers) and gene body (bottom, G12D allele methylated, WT methylated primers) were used on primary tumors (Primary/PT1) and DTCs. *In vitro* methylation with -/+ M.SssI was used as a control for primers. **H)** Overall survival (top) and curves showing accelerated cell death in the model when immunodeficient nude mice are used.

We next examined oncogenic pathways in PC such as the RAS pathway and its downstream targets, including PI3K, AKT, and Myc using the ultralow RNAseq dataset, and interestingly all were significantly down in the dormant DTC samples in comparison to the primary tumor or reactivated clones, (Fig. 6D).

The Ink4a.1 cell line is heterozygous for mutant KRAS, and yet RAS pathway targets were downregulated in dormant cells; therefore, we speculated that this could be due to selection of cells that suppressed the mutant allele. When we examined the single cell RNA-seq data at the KRAS locus, we observed that in the primary tumor cells there was expression of the KRAS^G12D^ allele; however; in two separate dormant samples almost all expressed KRAS was WT (Fig. 6E). Analysis of the *Cdkn2a* locus showed loss of expression of p16*Ink4a*, confirming the primary tumor and dormant samples were derived from the Ink4a.1 cell line (Fig. S12A).

To confirm these results, we then sorted DTCs from the livers of two dormant mice and harvested RNA for qPCR for mutant and WT KRAS using the Ink4a.1 cells as a control. We found equal expression of mutant and WT KRAS in Ink4a.1 cells, although in the dormant samples, only the WT allele was expressed (Fig. 6F). Gel analysis of the qPCR confirmed expression of the WT allele in all the samples whereas the mutant allele was detected in the Ink4a.1 cells only, with loss of expression in the dormant samples (Fig. S12B). We hypothesized that this suppression of mutant KRAS might be due to promoter methylation. Indeed, methylation-specific PCR for the CpG island located in the promoter of KRAS (Fig. S12C) revealed DNA methylation only in dormant DTCs, but not in the primary tumor (Fig. 6G upper panel). Since our parental Ink4a.1 cells were derived from a murine model containing the lox-stop-lox KRAS^G12D^ allele, we designed primers specific to the loxP site as a method to further confirm allele-specific silencing of the mutant KRAS allele using methylation-specific PCR of the KRAS gene body in the non-promoter region of the intron 2 (see Fig. S12C). Methylation of this region is associated with increased transcription (*44*) (*45*), and we found that this region was methylated in samples of sorted primary tumor cells and unmethylated in the dormant DTCs (Fig. 6G lower panel). Taken together we conclude this is evidence for monoallelic suppression of the mutant KRAS allele in dormant PC cells. This type of silencing of mutant KRAS is novel in cancer.

Several recent studies have implicated the innate and adaptive immune system in the regulation of dormancy, raising the question of whether dormancy is a cell autonomous or an extracellular driven process (*8*) (*46*). Pommier et. al. found in a PC model of dormant DTCs that unresolved ER stress lead to downregulation of MHCI as a mechanism of immune evasion (*46*). When we examined our single cell RNA-seq data for the expression of genes involved in ER stress we found that as a whole, this pathway was not significantly upregulated, although there were some samples with limited gene upregulation (i.e. Hspa5 (BiP) and EIF2AK3 (PERK), suggesting some cells are undergoing ER stress (Fig. S13A). Interestingly MHCI was not shown to be significantly differentially expressed between the dormant DTCs and primary tumor (data not shown). To determine what the expression would be for MHCI, we used flow cytometry of FACS-sorted dormant DTCs in comparison with the parental Ink4a.1 cells and found that MHCI was similarly expressed (not downregulated) (Fig. S13B). Nonetheless, we hypothesized that an immune mechanism likely contributed to dormancy in our model given our pathway analyses of the dormant transcriptome revealing a number of immunomodulators, checkpoint inhibitors, cytokines, and chemokines (i.e. Ccl5). To test this hypothesis, we performed the experiment using nude mice that were T cell-deficient and B cell-dysfunctional. We injected the same number of Ink4a.1 cells orthotopically into the pancreas, allowed primary tumors to form, and then resected them. The median survival of the immunodeficient group was 24 days versus 568 days for the immunocompetent group (Fig. 6H top panel). Strikingly we found that unlike the immunocompetent FVB background in which we would consistently observe 30-35% of the mice with the dormant phenotype, the immunocompromised nude mice had 100% die from early recurrence at a rate similar to the early recurrent phenotypic group in the FVB background (Fig. 6H bottom panel). This result indicates that the adaptive immune system is required for dormancy in this model. Taken together as whole, this set of data indicate that dormancy in this model is governed by a multitude of mechanisms that are both cell autonomous and non-autonomous.

## Discussion

We have produced the first mouse model of dormancy following resection of pancreatic cancer. Currently there are no models that reflect the biology of early stage resected pancreatic cancer, thus this model will be useful to conduct pre-clinical experiments testing of novel neoadjuvant and adjuvant therapeutics in PC. Our model provides the ability to study the underlying differences between early and latent recurrence, which our data suggest are different states, as well as test various therapeutic strategies to combat dormancy. One current controversy in the dormancy field relates to two very different therapeutic strategies to target dormancy. One focuses on pharmacologic agents that will drive non-dormant cells into dormancy (the “maintaining dormancy for life” strategy). The other focuses on pharmacologic agents that will “wake the cells up” so that they can be eradicated with other therapeutics (*47*). These important questions are difficult to resolve pre-clinically due to a lack of available models but could be tested in this model.

Taken together, our transcriptomic data highlight a multifaceted network of pathways and genes needed to maintain a state of pancreatic cancer dormancy (Figure 7). While some of these pathways have previously been shown to be important for dormancy in other systems, it is possible that others may be unique to either PC dormancy or dormancy in the liver. These include calcium signaling, neurotropic receptors (a possible explanation of perineural invasion that occurs in PC metastasis), and apolipoprotein expression, which are genes often expressed in the liver. For example, we identified Ccl5 as a gene highly expressed in murine dormant tumor cells from the liver and also highly expressed in human DTCs from the liver. The role of Ccl5 in cancer has largely been described in tumor progression and metastasis; however, there is some data supporting a role in immunosuppression through recruitment of T-regulatory lymphocytes (*39*). Whether that is the role that Ccl5 plays in PC dormancy remains to be seen, but nonetheless, Ccl5 appears to be a novel biomarker of PC dormancy.

**Figure 7.**
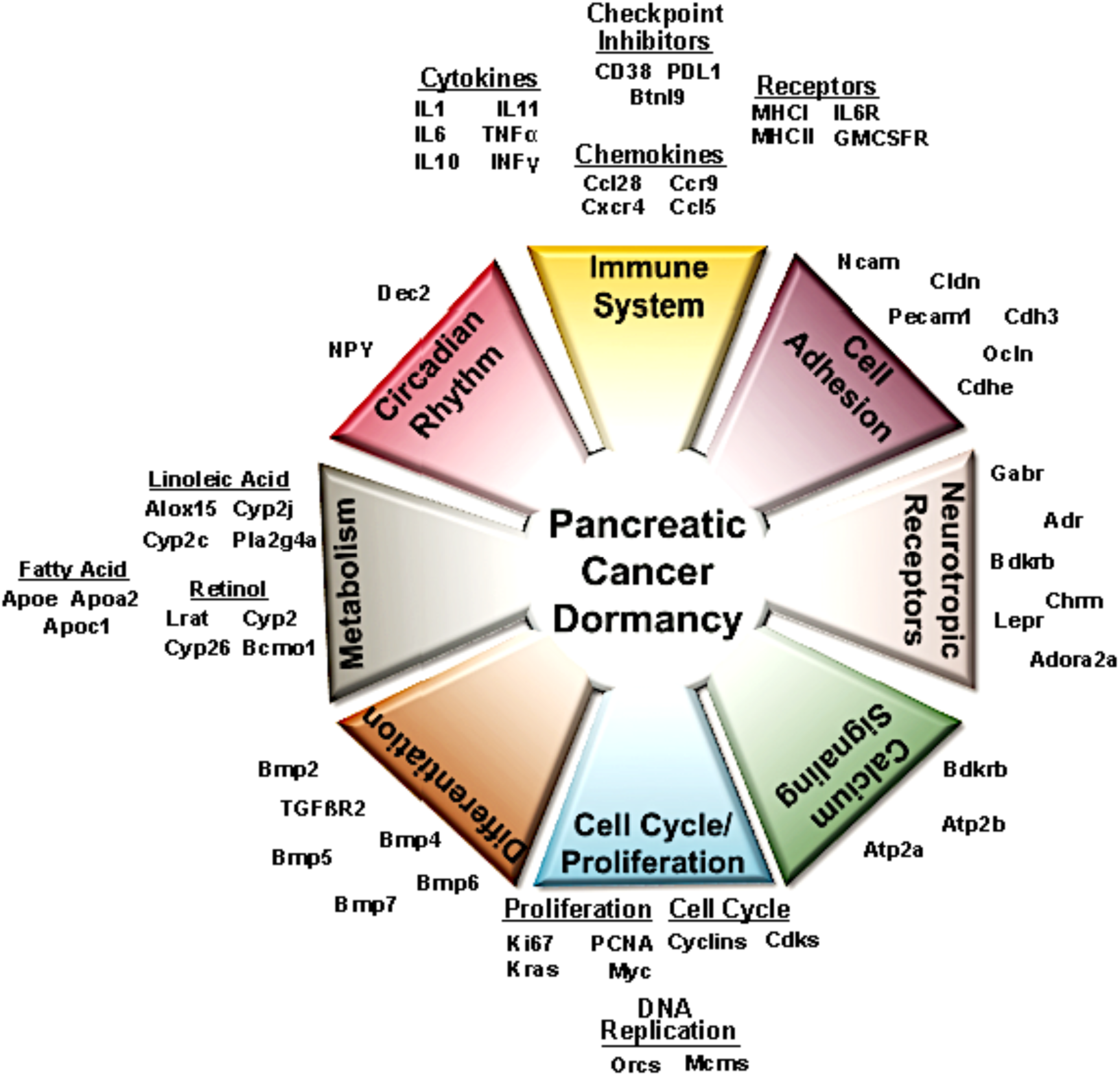
Model for pathways and genes needed for metastatic pancreatic cancer dormancy following resection of the primary tumor. Summary of data taken from expression and pathway analysis showing the complexity of intracellular mechanisms of metastatic pancreatic cancer cell dormancy. Note Cell Cycle/Proliferation pathway (blue wedge) is the only downregulated pathway homologous to both datasets.

One of the most significant findings in this study is the identification of a dormancy gene signature that has been validated in human disseminated tumor cells from the livers of patients undergoing surgery for localized pancreatic cancer. These are the first such studies of human disseminated tumor cells in patients with resectable pancreatic cancer and indicate that the livers of these patients contain a large abundance of DTCs. The pathways that are regulated by the genes in the signature have relevance to the biology of these DTCs in humans. Clearly PC patients fall into two groups after surgical resection: those that succumb to their disease after only a short time and those that experience a much better outcome, suggesting distinct disease trajectories. Yet what they share in common is the presence of DTCs at the time of the surgery of their primary tumor. These two outcomes cannot be explained by the sheer number of DTCs in the liver, which at the time of pancreatectomy is greater in the latent group then in the early recurrent group. We speculate that the fraction of DTCs that express the dormancy signature may correlate with a more latent or long-term survival phenotype. Perhaps this question can be answered not only from the liver DTC population but also from the blood using circulating tumors cells, which would be clinically more easily translatable.

Emerging mechanistic data has shown that a number of transcriptional modulators regulate dormancy, including NR2F1, DEC2, TGFβ2, and SOX9, which further supports the hypothesis that dormancy, like metastasis, is manifested by a state of cellular transdifferentiation that is epigenetically driven (*2*) (*28*) (*28*). There are numerous levels of evidence in our model that support the concept that dormancy reflects a change in cell state governed by epigenetic mechanism(s): 1) transcriptomic analysis supports a global change in dormancy that can change back to a non-dormant state, 2) expression of stem-cell like markers that are downregulated as cells come out of dormancy, 3) suppression of oncogenic signaling through monoallelic suppression of mutant KRAS, and 4) upregulation of transcriptional repressors like Dec2 that functionally regulate quiescence. The finding that the selective suppression of mutant KRAS expression through DNA methylation is novel in cancer and suggests an exquisite mechanism of DNA methylation reminiscent of imprinted methylation. Whether Dec2 is a master regulator of a greater number of pathways that are functionally relevant in dormancy remains to be seen. Our model provides a unique platform for us to study the epigenetic regulation of dormancy.

The process of epithelial to mesenchymal transition (EMT) in cancer produces cells with stem cell properties (*48*). It has been hypothesized that dormant tumor cells might share similar mechanisms of maintaining quiescence as stem cells (*48*). There are several lines of evidence that the dormant DTCs we isolated have properties of cancer stem cells. Increased expression of cell surface and cytosolic enzymes have been associated with pancreatic CSCs and increased expression of chemotherapy resistance genes (that are drug transporters and nucleotide metabolism genes) led to chemoresistant DTCs. Our data also identified several molecular pathways upregulated that are involved in maintaining other types of stem cell quiescence, such as BMPs. This illustrates how the mechanisms that maintain stem cell quiescence may be coopted by dormant cancer cells, as BMPs have been implicated in maintaining dormancy in a murine metastatic lung cancer model (*49*). It is possible that dormant cells are a quasi-stem cell that takes its cues/signals from the tissue in which it resides (*49*).

While we observed that a dormant tumor cell placed in culture can eventually “awaken” after several months of quiescence, we do not think this necessarily means that reactivation in this model is a cell autonomous process. Our data using nude mice as hosts indicate that the adaptive immune system is required for dormancy; whether it is needed to induce or just maintain dormancy is not clear at this point. This role of the immune system in our model is distinct from that of Pommier et. al. in which dormancy is characterized by a state of immune evasion (*8*). To the contrary, our data suggests that the adaptive immune system is required for dormancy and functions to hold these dormant DTCs in check. This result is more consistent with observations of awakening of dormant tumor cells in solid organ transplantation upon pharmacologic immunosuppression. This can occur in the transplanted organ when the donor or the recipient has a remote history of cancer (*50*). In both situations, upon application of immunosuppressive medicines to prevent allograft rejection, the awakening of dormant tumor cells is observed.

Another important question raised by this study is when do cancer cells adopt these transcriptomic changes? Does this happen during primary tumorigenesis or is this adapted after dissemination? Our data comparing the dormancy signature in mice to human resected pancreatic primary tumors suggests that perhaps some of these changes may occur before dissemination. Nonetheless, if cells in the primary tumor do adopt a dormancy phenotype then perhaps this dormancy signature can be used to predict outcome (early recurrent versus latent) after surgery on circulating tumor cells. This potentially could guide clinical management decisions when often elderly patients are being considered for a highly morbid operation.

In conclusion, we have developed a new model of pancreatic cancer dormancy that has provided several mechanistic insights into the biology of this rare cell population. Most notably cancer dormancy is governed by multiple cell autonomous and non-cell autonomous mechanisms working in concert. Whether these mechanisms can all be explained by epigenetic superenhancers needs to be investigated. Nonetheless, these changes allow the cell to adopt stem-like properties and to evade key oncogenic signaling pathways that normally drive tumorigenesis. Importantly, these changes are highly plastic as cells can come out of dormancy and revert back to their pre-dormant cellular programs. This suggests that understanding these programs should lead to the ability to manipulate dormancy therapeutically. If dormant tumor cells are phenotypically similar to quiescent stem cells, then perhaps strategies to drive them into senescence would be attractive.

## Materials and Methods

### Animal studies

All animal protocols were approved by the Rutgers Biomedical and Health Sciences Animal Care and Use Committee. Five-to six-week old female FVB mice were purchased from The Jackson Laboratory (Bar Harbor, ME). For orthotopic studies, 100 Ink4a.1 luc/mcherry pancreatic cells in a mixture of 50% matrigel/50% DMEM +10% FBS were injected orthotopically into the tail of the pancreas. Mice were imaged via IVIS imaging. Images were set at the same radiance scale using LivingImage software. For distal pancreatectomy with splenectomy surgery, an incision was made in the abdomen to expose the primary pancreatic tumor and spleen. The lower gastric vessel was clamped and cut proximal to the spleen. The healthy pancreas was clamped and cut. The primary tumor and spleen were removed en mass. The abdominal wall was closed with 5-0 vicryl sutures and skin closed with wound clips. Buprenorphine and bupivacaine were given for pain management and the mouse was allowed to recover under a heating lamp until ambulatory. Mice were monitored for signs of pancreatic cancer recurrence twice a week. For intrasplenic injections, 10 to 1 million cells were injected into the spleen near the hilum. The spleen was removed as above except that a medium clip was placed across the tail of the pancreas and cut proximal to the spleen.

### Ultralow and standard RNA-seq analysis

The disseminated cells were profiled using ultralow input RNA-seq protocol while primary tumor and metastatic cells were profiled using standard RNA-seq.After quality check of reads using FastQC (http://www.bioinformatics.babraham.ac.uk/projects/fastqc), we used Salmon (*50*) to quantify transcript-level expression and EdgeR (*51*) to identify genes with significantly differential expression between pairs of conditions based on replicated count data from bulk RNA-seq profiling. The normalized data were applied to R package GAGE (*52*) for gene-set enrichment and pathway analysis. The p-values were corrected for multiple testing using FDR. Pathview (*52*) was used to identify and visualize KEGG pathways significantly enriched for differentially expressed genes. Heat maps were created using Morpheus from the Broad Institute (https://software.broadinstitute.org/morpheus).

## Acknowledgements

We would like to thank Paul Labhart from Adaptive Motif for analysis of ATAC-seq data. This work was supported by grants from the National Cancer Institute (K08 CA172676, R01 CA200800, D.R.C.), Breast Cancer Research Foundation (D.R.C.), Pancreatic Cancer Action Network Translational Research Grant (D.R.C and C.D.), National Institute of General Medical Sciences (R01GM129066, S.D.), and New Jersey Alliance for Clinical and Translational Science (B.G.). This was also funded in part by a donation from the Eleanor Russo fund for Pancreatic Cancer Research. We thank Rutgers Cancer Institute of New Jersey Histology core facility for processing the tumor blocks. Flow cytometry research was generated by the Rutgers Cancer Institute of New Jersey Flow Cytometry and Cell Sorting Shared Resource, supported, in part, with funding from NCI-CCSG P30CA072720-5921.

## Notes

### Competing Interest Statement

The authors have declared no competing interest.

