## Supplemental Materials for "A Novel Model of Pancreatic Cancer Dormancy Reveals Mechanistic Insights and a Dormancy Gene Signature with Human Relevance"

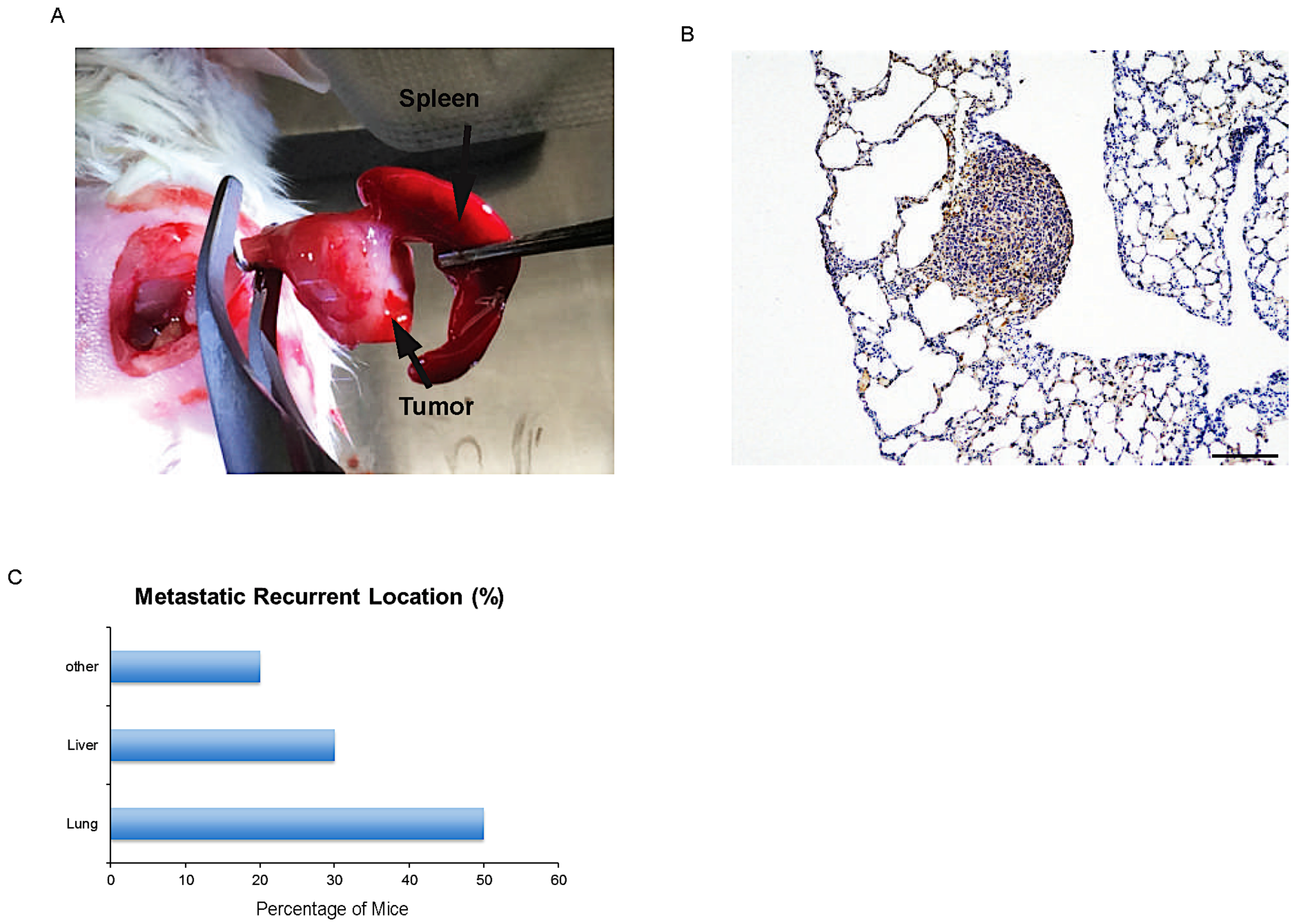


**Supplemental Figure 1. An orthotopic syngeneic resectable model of metastatic pancreatic cancer. A.** Pancreatectomy with splenectomy. After cauterization of the lower gastric vessel, the spleen and primary pancreatic tumor are removed en mass by bisecting the pancreas with a medium clip and excising proximal to the tumor. Arrows showing location of spleen and primary pancreatic tumor. **B.** A latent (DFI>200 days) pancreatic cancer metastasis expressing mCherry in the lung. Scale bar: 200 µm. **C.** Location of recurrences in latent recurrent mice. n=10. The majority of metastasis arises in the lung and liver.

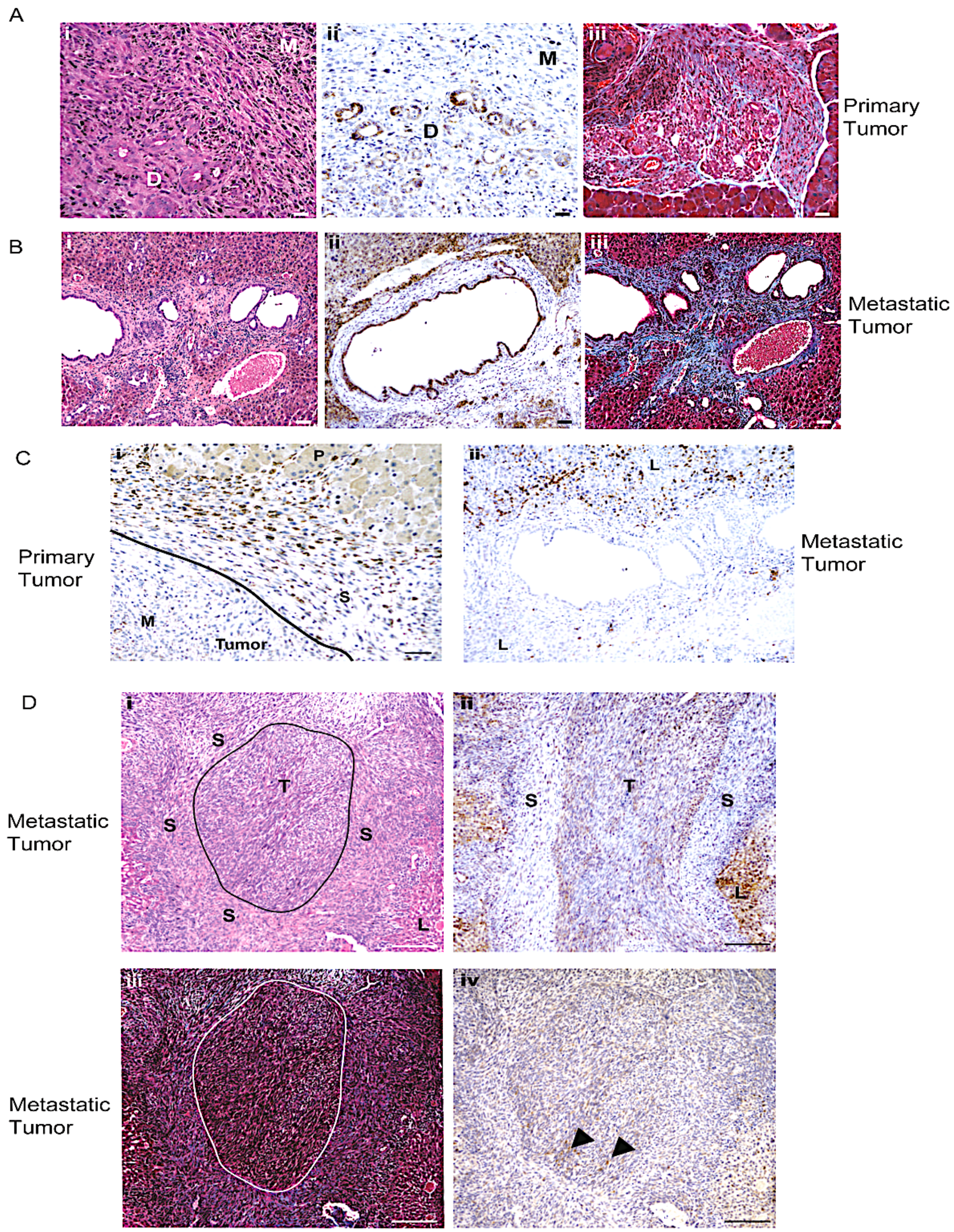


**Supplemental Figure 2. CSC-like primary tumor cells give rise to well-differentiated and quasi-mesenchymal liver metastasis indicative of chromatin remodeling. A.** Histology of a representative primary pancreatic tumor resected at day 28 post-cell injection. **(i)** H&E, **(ii)** E-cadherin staining displaying loss of staining in mesenchymal-like cells, with darker staining present in well-differentiated cells. **(iii)** Masson's trichrome stain showing collagen deposition in blue. Notice the increased concentration of collagen along the periphery of the tumor. Scale bar, 50 µm. **B.** Histology of a representative metastatic tumor to the liver displaying well-differentiated features. **(i)** H&E, **(ii)** E-cadherin, **(iii)** Masson's Trichrome. Scale bar, 50 µm. D: well-differentiated tumor cells, M: mesenchymal-like cells. **C.** CD68 expression in a primary tumor (left) and glandular-like PC liver metastasis (right). M: mesenchymal cells, S: stroma, P: pancreas, L: liver. Scale bar (left): 50 µm, (right): 20 µm **D.** Histology of a representative metastatic pancreatic tumor to the liver displaying mesenchymal features. **(i)** H&E with tumor border drawn, **(ii)** E-cadherin, **(iii)** Masson's trichrome stain, with tumor border drawn in white, **(iv)** CD68 stain. Arrowheads point to $CD68^{+}$ cells along tumor edge. Scale bar, 50 µm. S: stroma, T: tumor.

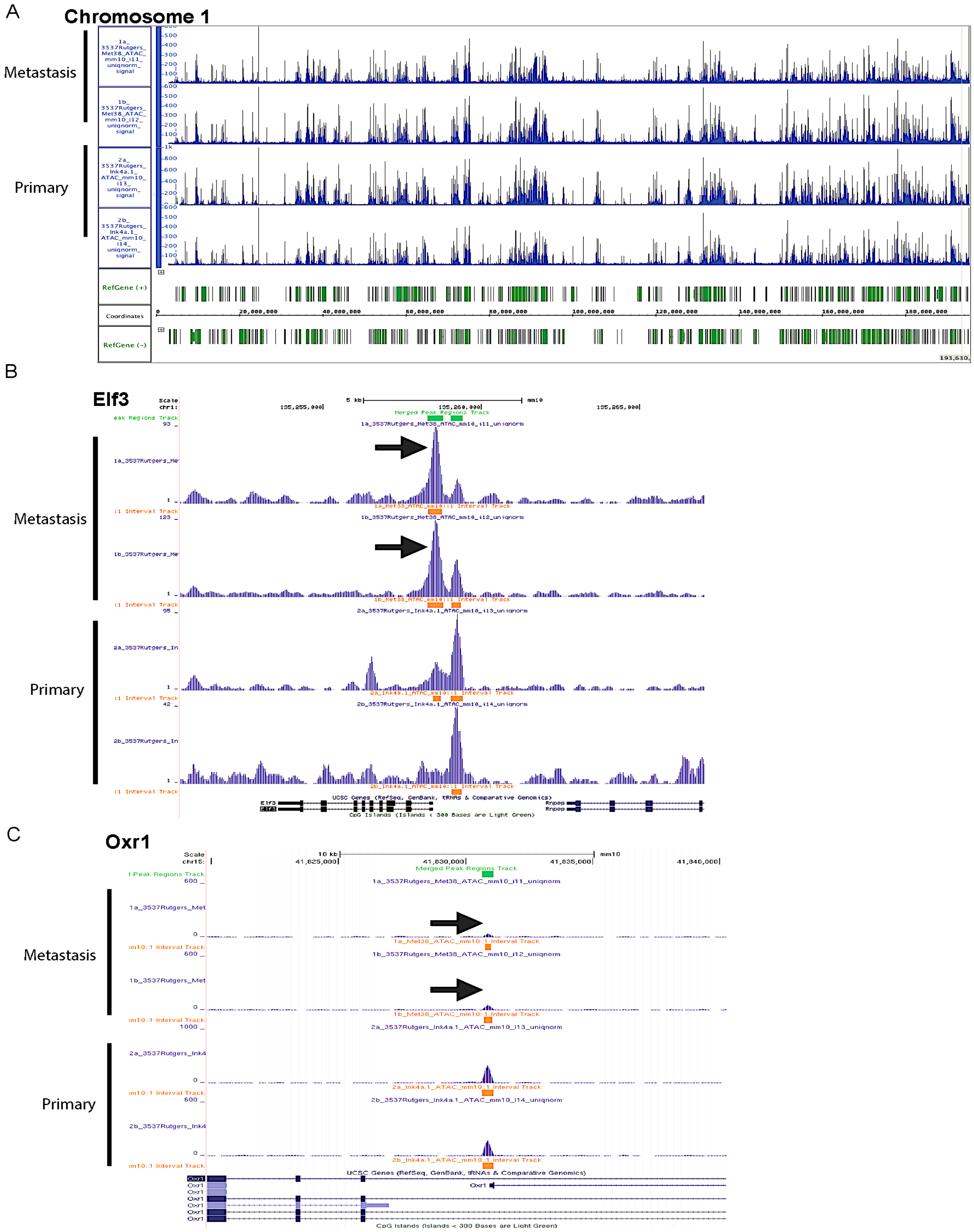


**Supplemental Figure 3. ATAQ-seq data displays variations in open chromatin between primary and metastatic pancreatic cancer. A.** Example of open regions (blue peaks) of chromosome 1 in the Ink4a.1 (Primary) cell line versus the early metastatic cell line Met38 (Metastasis) using the Integrated Genome Browser (Bioviz.org). Reference genes are in green, genomic coordinates are in black, n=2 each cell line. **B.** *Elf3* locus showing an increase (arrow) in the accessibility at the promoter in the metastatic tumor. Schematic of *Elf3* gene structure is in black on the bottom of the figure. Merged peak regions are in green at the top of the figure. **C.** *Oxr1* locus showing a decrease (arrow) in the accessibility at the promoter in the metastatic tumor. Schematic of *Oxr1* gene structure is in black on the bottom of the figure. Merged peak regions are in green at the top of the figure.

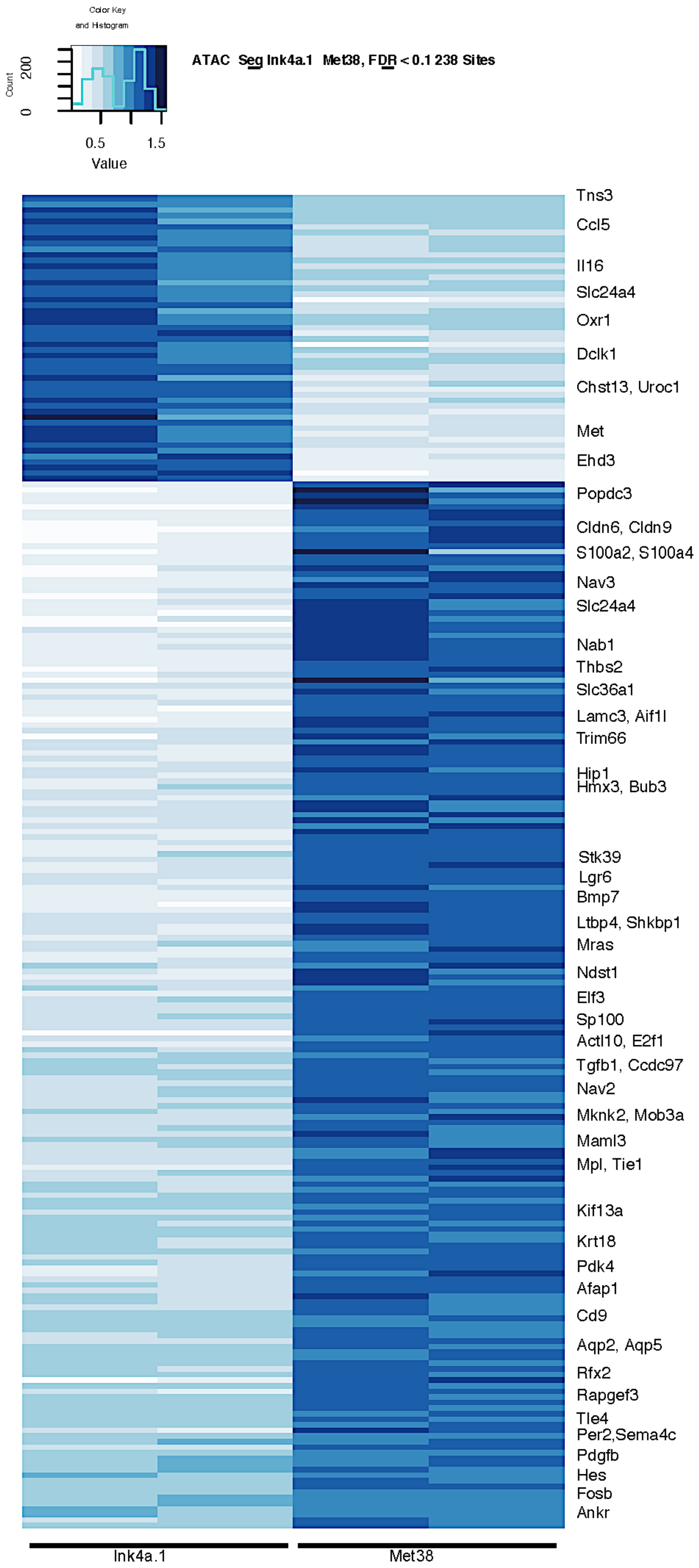


**Supplemental Figure 4. Differential changes in the accessibility of chromatin from metastatic tumors.** ATAQ-seq analysis of primary tumor cell line, Ink4a.1, in comparison with metastatic pancreatic cell line, Met38. n=2 each cell line. Dark blue denotes regions of open areas of chromatin. A partial listing of genes within the euchromatin are listed to the right.

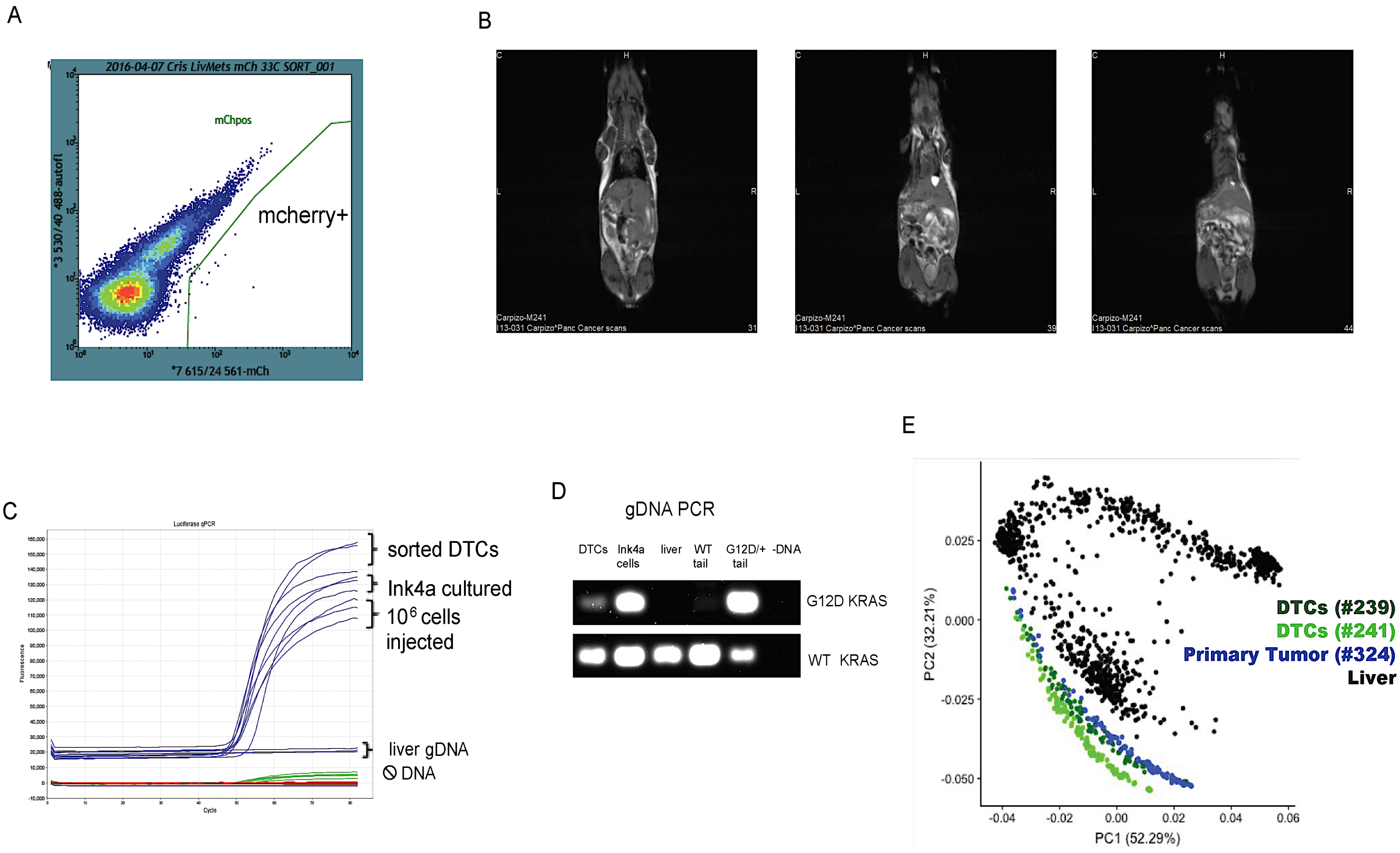


**Supplemental Figure 5. Metastatic DTCs from the livers of latent recurrent dormant mice are pancreatic cancer cells. A.** mCherry$^{+}$ cells were sorted from the liver of a mouse with DFI>300 days. **B.** Full body MRI scans of mouse #241 used for single cell RNA sequencing showed no evidence of gross disease. **C.** Presence/absence protocol was used for Taqman luciferase PCR on genomic DNA. Genomic DNA from cultured Ink4a.1 cells and metastatic liver was used as positive controls. WT liver genomic DNA and no DNA were used as negative controls. **D.** Mutant-specific PCR for Kras$^{G12D}$ and wild-type Kras was completed on genomic DNA from the same mouse as in A. Genomic DNA from cultured Ink4a.1 cells and tail DNA from a KPC mouse were used as positive controls. WT liver and tail DNA were used as positive controls for WT Kras PCR and negative controls for mutant Kras PCR and no DNA was used as a negative control for both. **E.** Principle component analysis displaying the distribution of cells based on their expression (taken from the 10x genomics analysis). Samples include the primary tumor (dark blue), two dormant DTC samples (light and dark green), and liver cells (black), which are a source of possible contamination.

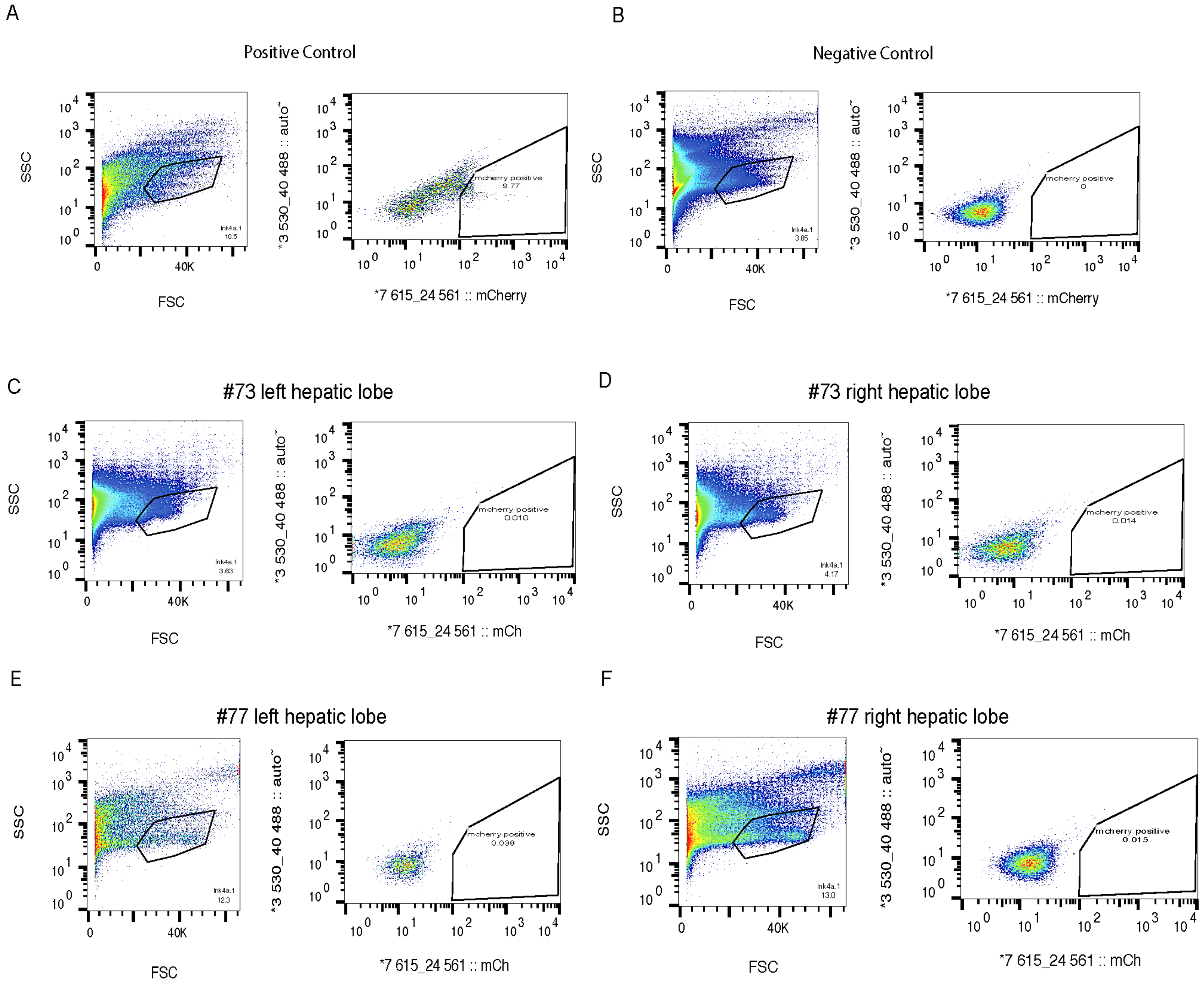


**Supplemental Figure 6. Metastatic DTCs are found in the livers of mice at the time of primary tumor removal.** Liver single cell suspensions were made either from mice splenically-injected with Ink4a.1 (**A**, positive control), PBS (**B**, negative control), or orthotopically-injected mice with a primary tumor present. Flow cytometry from suspensions of the left hepatic lobe (**C**) and right hepatic lobe (**D**) of mouse #73 was used to find mCherry$^{+}$ cells. **E.** Flow cytometry of the left hepatic lobe and right hepatic lobe **(F)** of mouse #77 completed at the time of primary tumor removal.

A

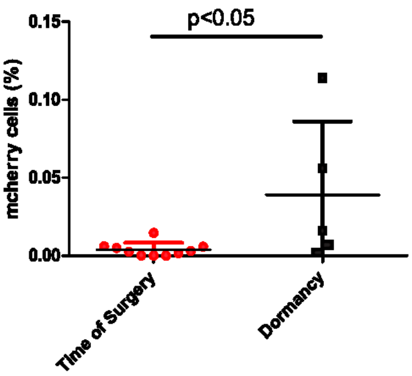


B

Organ dissemination frequency (%)

| Organ | Mouse Number | | |
|---|---|---|---|
| | 239 | | 249 |
| bone marrow | 0.083 | 0.002 | 0.001 |
| brain | 0 | 0.004 | 0.016 |
| heart | 0.019 | 0.012 | 0.078 |
| intestine | 0 | 0 | 0.462 |
| kidney | 0.017 | 0 | 0.036 |
| liver | 0.294 | 0.027 | 0.001 |
| lung | 0.005 | 0 | 0 |
| stomach | 0.620 | 0.204 | 0 |

**Supplemental Figure 7. Metastatic PC dormant cells are chemoresistant and express markers of CSCs. A.** Frequency of mCherry+ cells in the liver of mice at the time of surgery (average DFI=0, n=10) and during the dormancy period (average DFI=262 days, n=5). **B.** Organ dissemination frequency of mCherry+ cells in organs listed, n=3. Average DFI=165 days.

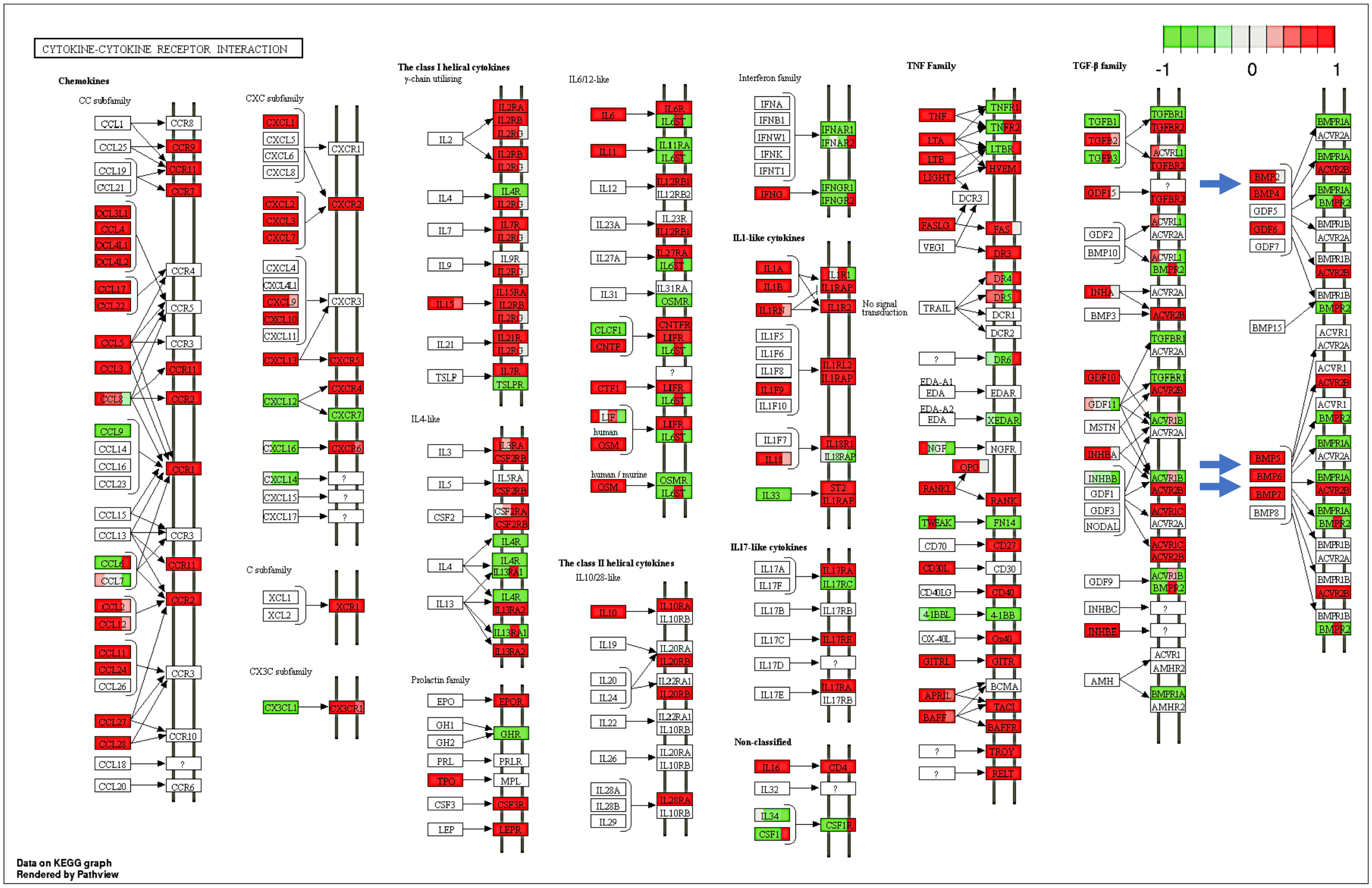


**Supplemental Figure 8. The cytokine-cytokine receptor interaction pathway is enriched with differentially upregulated gene sets.** The majority of genes in the cytokine-cytokine receptor KEGG pathway (mmu04060) were upregulated. Red boxes denote upregulation of the labeled gene (or protein) while green denotes downregulation. Boxes that are multicolored represent isoform expression of that particular gene. Arrows are highlighting the overexpression of BMPs. Pathway analysis results were visualized using R package Pathview and KEGG pathways.

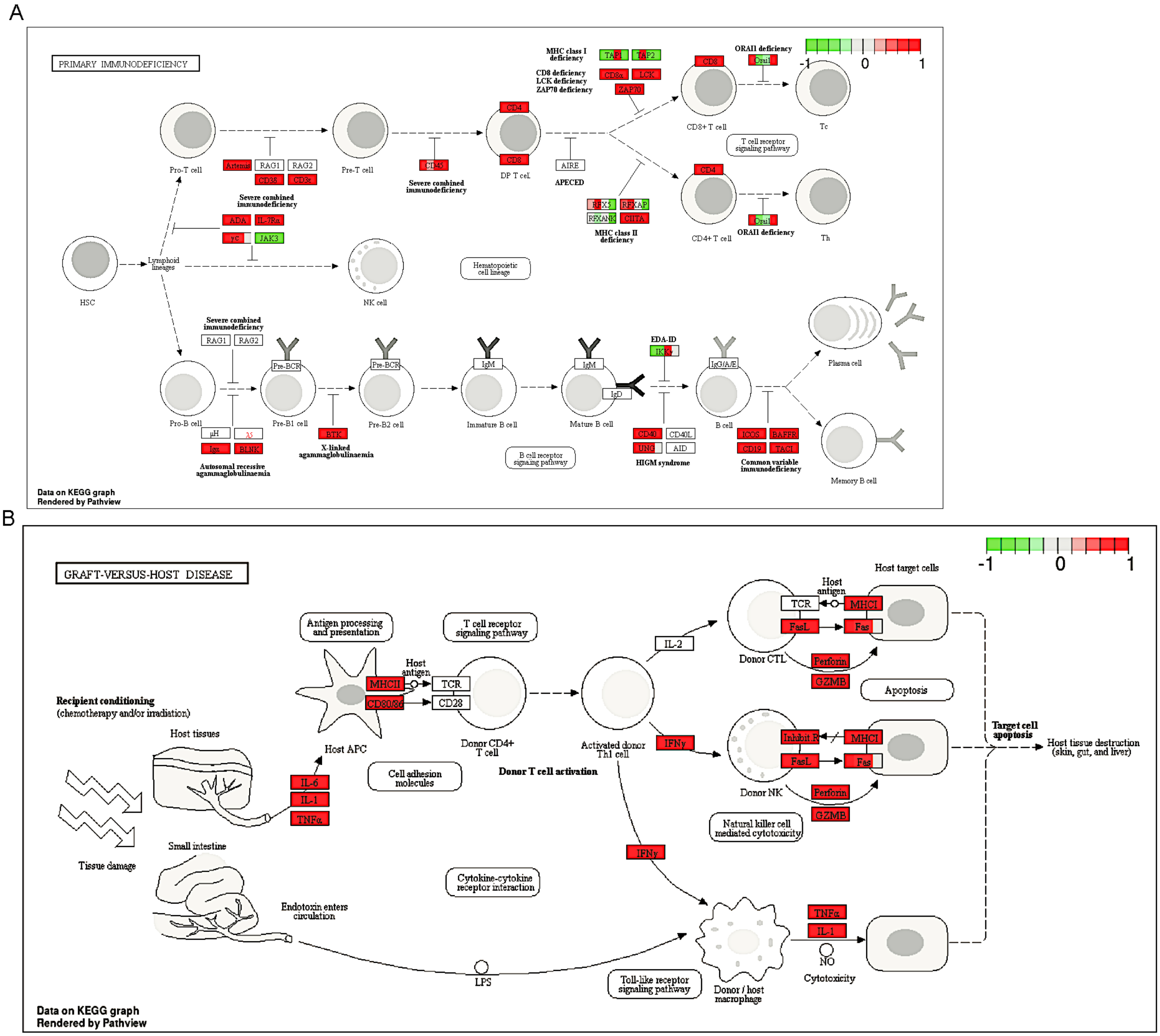


**Supplemental Figure 9. Immunological pathways are enriched with differentially upregulated gene sets in both ultralow and 10x genomic single cell expression datasets.** **A**. Primary immunodeficiency (mmu05340) and (**B)** graft vs host disease (mmu05332) KEGG pathways. Red boxes denote upregulation of the labeled gene (or protein) while green denotes downregulation. Boxes that are multicolored represent isoform expression of that particular gene. Pathway analysis results were visualized using R package Pathview and KEGG pathways.

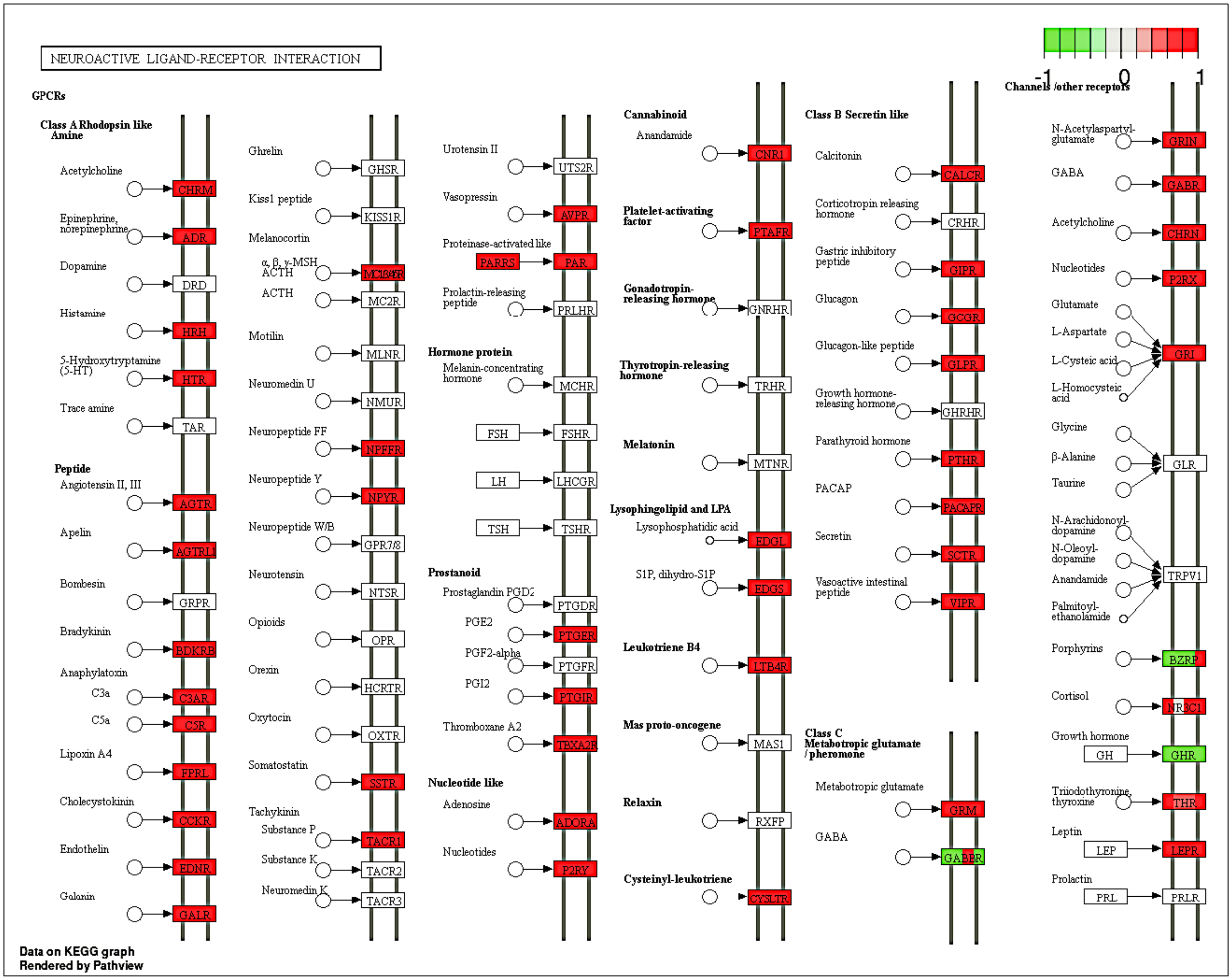


**Supplemental Figure 10. Neuroactive ligand-receptor pathways are enriched with differentially upregulated gene sets in both ultralow and 10x genomic single cell expression datasets.** Most receptors in the KEGG pathway for neuroactive ligand-receptor interaction (mmu04080) are upregulated in dormant pancreatic cancer cells. Red boxes denote upregulation of the labeled gene (or protein) while green denotes downregulation. Boxes that are multicolored represent isoform expression of that particular gene. Pathway analysis results were visualized using R package Pathview and KEGG pathways.

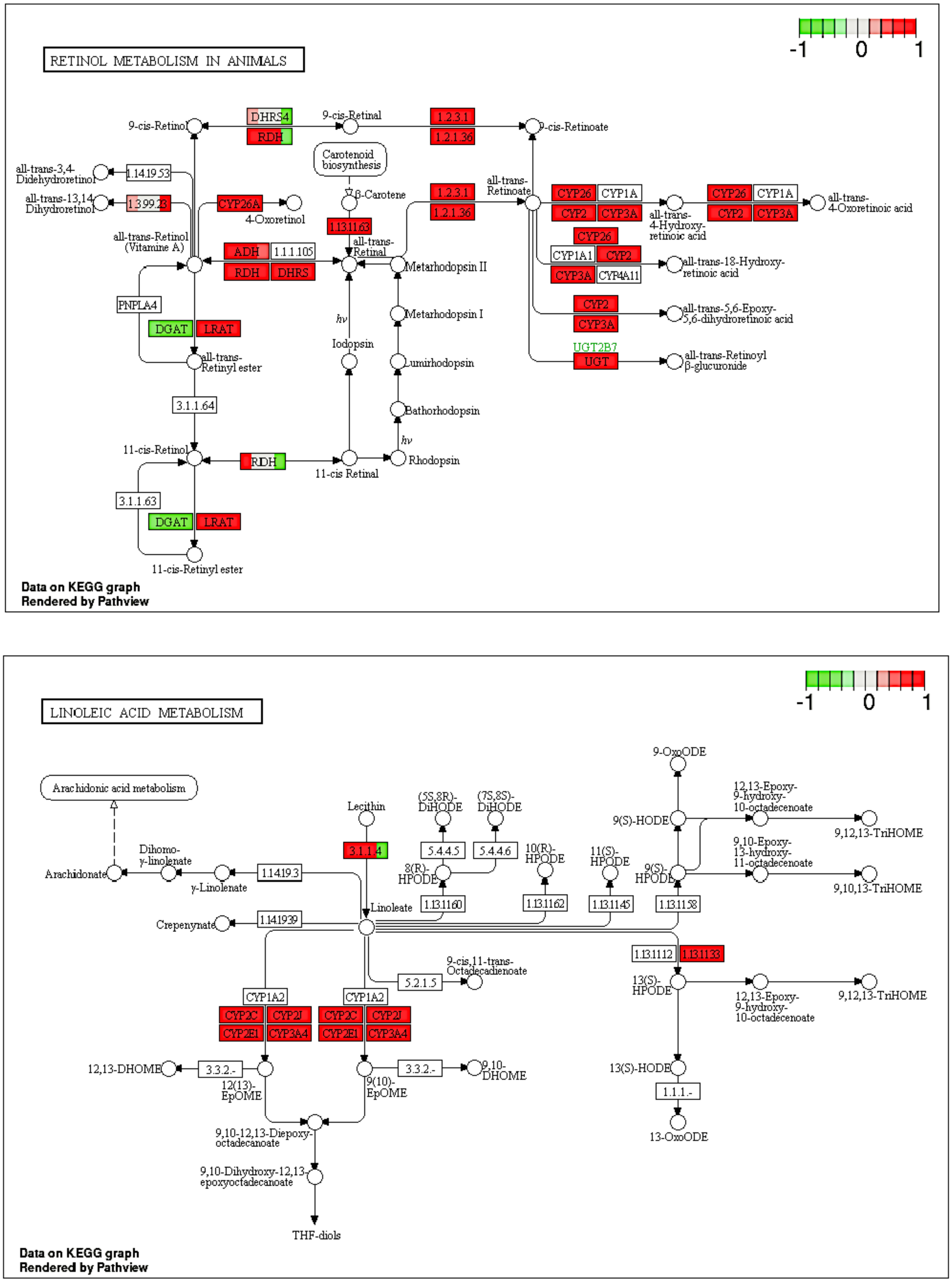


**Supplemental Figure 11. Metabolic pathways are enriched with differentially upregulated gene sets in both ultralow and 10x genomic single cell expression datasets.** **A**. Genes upregulated in the KEGG pathway for retinol metabolism (mmu00830) and **(B)** linoleic metabolism (mmu00591) suggests a possible dependence to maintain dormancy. Red boxes denote upregulation of the labeled gene (or protein) while green denotes downregulation. Boxes that are multicolored represent isoform expression of that particular gene. Pathway analysis results were visualized using R package Pathview and KEGG pathways.

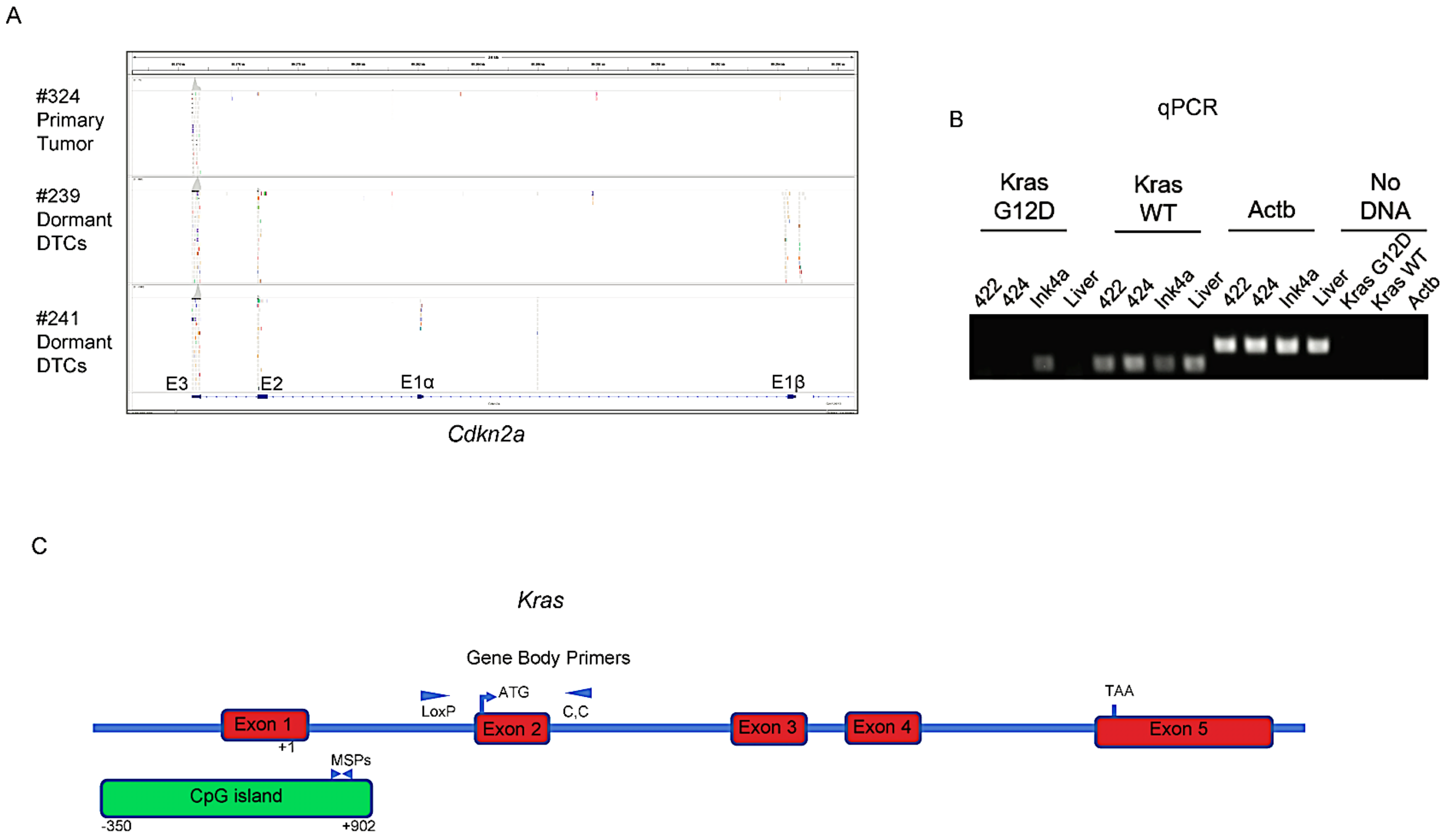


**Supplemental Figure 12. Dormant DTCs have lost expression of mutant *Kras*. A.** IGV analysis of expression of *Cdkn2a* in the primary tumor #324, dormant DTCs #239, and dormant DTCs #241. Loss of Exon 1α (E1α) expression is indicative of loss of p16Ink4a. **B.** Agarose gel analysis of qPCR for KrasG12D, Kras WT, and actin used on cDNA taken from two samples of DTCs (422, 424), the Ink4a.1 cultured cells, and normal FVB liver. No DNA was used for the negative controls. **C.** Schematic drawing of the Kras gene locus. Methylation-specific PCR (MSP) primers are shown located in intron 1, within the CpG island. Gene Body Primers encompass the LoxP sequence in the Kras G12D allele and wildtype sequence of exon 2.

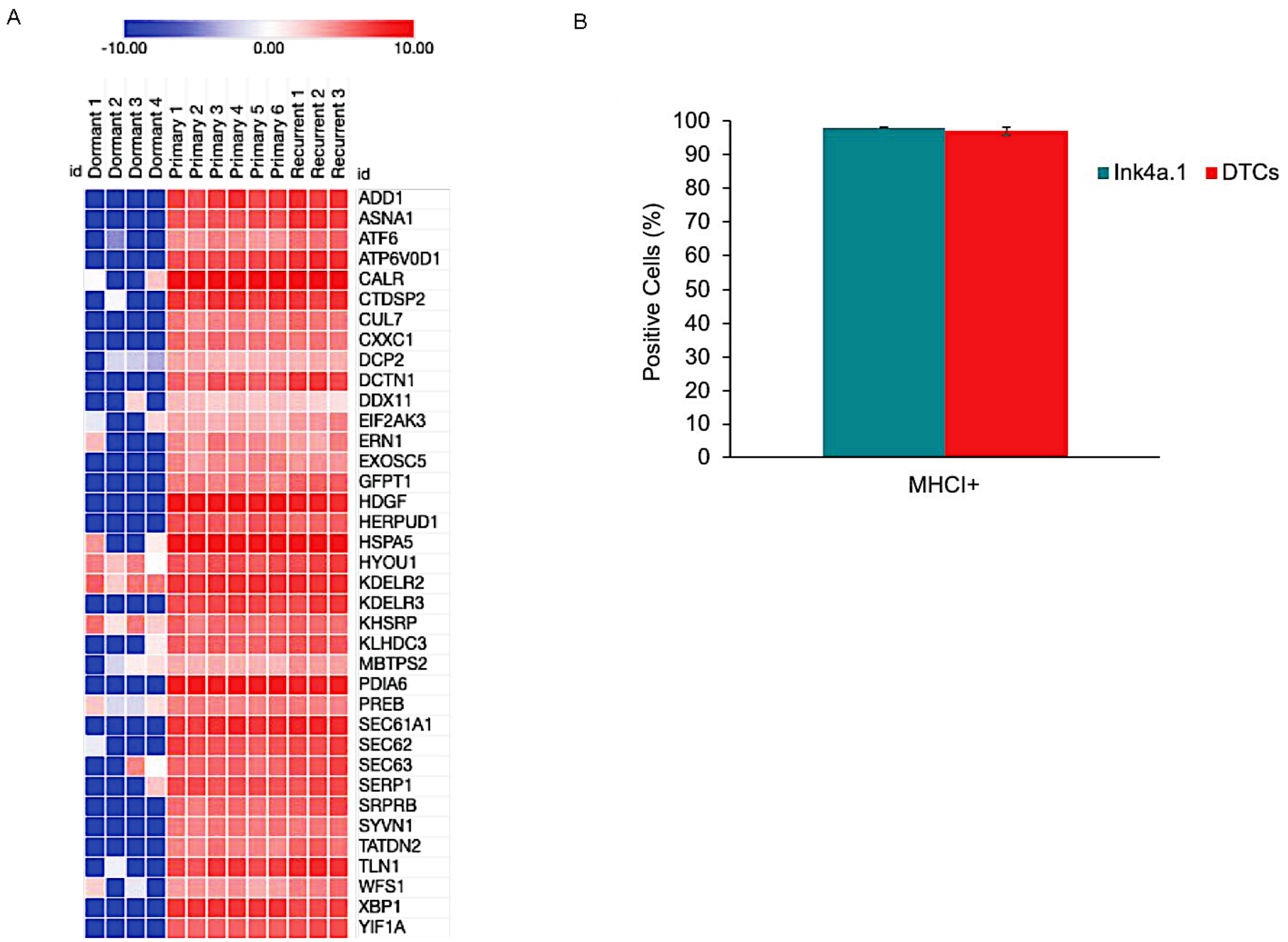


**Supplemental Figure 13. Dormant DTCs have mixed expression of ER Stress genes and express MHCI. A.** Expression of ER stress markers analyzed from ultralow RNA-seq analysis of individual dormant DTCs (dormant 1-4) compared with primary tumors (Primary 1-6), and *in vitro* recurrent cells (Recurrent 1-3). Transcript expression was calculated using the $\log_2$ values of the significant non-zero values of TPMs. Heat maps were created using Morpheus from the Broad Institute at https://software.broadinstitute.org/morpheus/ **B.** Flow cytometry analysis of DTCs stained for MHCI using a mouse H-2K$^d$/H-2D$^d$ MHC class I-APC antibody (clone 34-1-2S, detects q haplotype of FVB strain). Single cell suspensions from the livers of dormant mice were analyzed for expression of CD44, CD133, mCherry (DTC markers) and MHCI expression. The parental primary tumor cell line Ink4a.1 was used as a control.

Table S1. KEGG Pathways with Upregulated Genes in Dormant Single Cells

| Pathway Name | p value | q value |
|---|---|---|
| mmu05332 Graft-versus-host disease | 1.09E-14 | 2.21E-12 |
| mmu05330 Allograft rejection | 2.47E-13 | 2.49E-11 |
| mmu04610 Complement and coagulation cascades | 1.09E-11 | 6.87E-10 |
| mmu04612 Antigen processing and presentation | 1.36E-11 | 6.87E-10 |
| mmu05320 Autoimmune thyroid disease | 3.03E-11 | 1.23E-09 |
| mmu05150 Staphylococcus aureus infection | 9.10E-11 | 3.06E-09 |
| mmu04650 Natural killer cell mediated cytotoxicity | 1.47E-09 | 4.23E-08 |
| mmu04060 Cytokine-cytokine receptor interaction | 2.69E-09 | 6.79E-08 |
| mmu04640 Hematopoietic cell lineage | 5.37E-09 | 1.21E-07 |
| mmu05340 Primary immunodeficiency | 1.03E-08 | 2.09E-07 |
| mmu04660 T cell receptor signaling pathway | 2.13E-08 | 3.91E-07 |
| mmu04672 Intestinal immune network for IgA production | 4.52E-08 | 7.61E-07 |
| mmu04380 Osteoclast differentiation | 5.23E-08 | 8.13E-07 |
| mmu04940 Type I diabetes mellitus | 7.20E-08 | 1.04E-06 |
| mmu04514 Cell adhesion molecules (CAMs) | 1.13E-07 | 1.52E-06 |
| mmu05140 Leishmaniasis | 1.61E-07 | 2.04E-06 |
| mmu04145 Phagosome | 9.42E-07 | 1.12E-05 |
| mmu05416 Viral myocarditis | 1.56E-06 | 1.75E-05 |
| mmu05310 Asthma | 3.08E-06 | 3.28E-05 |
| mmu04062 Chemokine signaling pathway | 4.62E-06 | 4.66E-05 |
| mmu04662 B cell receptor signaling pathway | 1.04E-05 | 1.00E-04 |
| mmu05322 Systemic lupus erythematosus | 1.20E-05 | 1.10E-04 |
| mmu04740 Olfactory transduction | 2.60E-05 | 2.29E-04 |
| mmu05323 Rheumatoid arthritis | 2.83E-05 | 2.38E-04 |
| mmu04620 Toll-like receptor signaling pathway | 4.68E-05 | 3.78E-04 |
| mmu05142 Chagas disease (American trypanosomiasis) | 5.54E-05 | 4.30E-04 |
| mmu04621 NOD-like receptor signaling pathway | 6.77E-05 | 5.06E-04 |
| mmu04080 Neuroactive ligand-receptor interaction | 2.08E-04 | 1.50E-03 |
| mmu04630 Jak-STAT signaling pathway | 2.23E-04 | 1.55E-03 |
| mmu04670 Leukocyte transendothelial migration | 2.59E-04 | 1.69E-03 |
| mmu05020 Prion diseases | 2.59E-04 | 1.69E-03 |
| mmu00983 Drug metabolism - other enzymes | 4.53E-04 | 2.86E-03 |
| mmu05145 Toxoplasmosis | 6.73E-04 | 4.12E-03 |
| mmu05143 African trypanosomiasis | 1.16E-03 | 6.89E-03 |
| mmu00830 Retinol metabolism | 1.48E-03 | 8.55E-03 |
| mmu03320 PPAR signaling pathway | 1.74E-03 | 9.79E-03 |
| mmu04976 Bile secretion | 2.55E-03 | 1.39E-02 |
| mmu02010 ABC transporters | 3.66E-03 | 1.95E-02 |
| mmu04664 Fc epsilon RI signaling pathway | 5.31E-03 | 2.75E-02 |
| mmu00591 Linoleic acid metabolism | 1.04E-02 | 5.22E-02 |
| mmu04142 Lysosome | 1.06E-02 | 5.22E-02 |
| mmu04020 Calcium signaling pathway | 1.16E-02 | 5.56E-02 |
| mmu05144 Malaria | 1.63E-02 | 7.65E-02 |
| mmu04666 Fc gamma R-mediated phagocytosis | 2.04E-02 | 9.35E-02 |

Pathway analysis was completed as described in Methods on single cell samples from the 10X Genomics data. Only pathways with q<0.1 are displayed.

**Table S1. Genes located in differentially open and closed regions of dormant DTCs.** ATAQ-seq analysis of differentially open and closed regions of the chromatin in dormant DTCs when compared with a recurrent tumor. The DESeq2 algorithm was used, and genes are listed with a $\log_2$ fold change>2 or <-2 and with an FDR<0.1.

### Table S2. KEGG Pathways with Upregulated Genes in Dormant Cells from Ultralow Input Data

| Pathway Name | p value | q value |
|---|---|---|
| mmu04740 Olfactory transduction | 2.35E-126 | 4.80E-124 |
| mmu04080 Neuroactive ligand-receptor interaction | 1.38E-33 | 1.41E-31 |
| mmu04640 Hematopoietic cell lineage | 6.61E-06 | 4.13E-04 |
| mmu04060 Cytokine-cytokine receptor interaction | 8.09E-06 | 4.13E-04 |
| mmu04950 Maturity onset diabetes of the young | 2.90E-05 | 1.18E-03 |
| mmu05340 Primary immunodeficiency | 1.58E-04 | 5.38E-03 |
| mmu04020 Calcium signaling pathway | 3.50E-04 | 1.02E-02 |
| mmu00830 Retinol metabolism | 4.23E-04 | 1.08E-02 |
| mmu00140 Steroid hormone biosynthesis | 7.48E-04 | 1.70E-02 |
| mmu04514 Cell adhesion molecules (CAMs) | 8.60E-04 | 1.75E-02 |
| mmu04672 Intestinal immune network for IgA production | 9.64E-04 | 1.79E-02 |
| mmu05332 Graft-versus-host disease | 1.10E-03 | 1.87E-02 |
| mmu00910 Nitrogen metabolism | 2.07E-03 | 3.24E-02 |
| mmu00590 Arachidonic acid metabolism | 2.69E-03 | 3.92E-02 |
| mmu00591 Linoleic acid metabolism | 3.07E-03 | 4.18E-02 |
| mmu00982 Drug metabolism - cytochrome P450 | 3.72E-03 | 4.75E-02 |
| mmu05143 African trypanosomiasis | 6.04E-03 | 7.25E-02 |
| mmu00980 Metabolism of xenobiotics by cytochrome P450 | 7.06E-03 | 8.00E-02 |

Pathway analysis was completed as described in Methods on single cell samples from the ultralow RNAseq data. Only pathways with q<0.1 are displayed.

**Table S2. KEGG Pathways with Upregulated Genes in Dormant Single Cells.** 10X genomics RNA-seq analysis was carried out and analyzed as described in Methods. Pathway enrichment was completed using GAGE (see Methods for more detail). Pathways with upregulated genes present are listed with q<0.1.

Table S3. KEGG Pathways with Downregulated Genes in Dormant Single Cells

| Pathway Name | p value | q value |
|---|---|---|
| Single cell data | | |
| mmu04110 Cell cycle | 2.15E-04 | 4.34E-02 |
| Ultralow cell data | | |
| mmu03010 Ribosome | 9.00E-70 | 1.84E-67 |
| mmu04141 Protein processing in endoplasmic reticulum | 5.87E-33 | 5.98E-31 |
| mmu03040 Spliceosome | 1.17E-23 | 7.98E-22 |
| mmu05016 Huntington's disease | 1.92E-22 | 9.79E-21 |
| mmu05012 Parkinson's disease | 1.14E-21 | 4.64E-20 |
| mmu00190 Oxidative phosphorylation | 2.08E-21 | 7.09E-20 |
| mmu03013 RNA transport | 1.42E-16 | 4.13E-15 |
| mmu05010 Alzheimer's disease | 1.28E-13 | 3.27E-12 |
| mmu04142 Lysosome | 1.62E-13 | 3.67E-12 |
| mmu04144 Endocytosis | 6.34E-13 | 1.29E-11 |
| mmu04120 Ubiquitin mediated proteolysis | 9.13E-12 | 1.69E-10 |
| mmu04510 Focal adhesion | 2.18E-11 | 3.71E-10 |
| mmu03050 Proteasome | 3.16E-11 | 4.96E-10 |
| mmu05220 Chronic myeloid leukemia | 1.58E-09 | 2.31E-08 |
| mmu05212 Pancreatic cancer | 2.34E-09 | 3.19E-08 |
| mmu00020 Citrate cycle (TCA cycle) | 3.05E-09 | 3.89E-08 |
| mmu04110 Cell cycle | 5.50E-09 | 6.60E-08 |
| mmu00240 Pyrimidine metabolism | 1.05E-08 | 1.19E-07 |
| mmu03060 Protein export | 2.23E-08 | 2.39E-07 |
| mmu03015 mRNA surveillance pathway | 3.99E-08 | 4.07E-07 |
| mmu05100 Bacterial invasion of epithelial cells | 6.90E-08 | 6.70E-07 |
| mmu04145 Phagosome | 4.03E-07 | 3.74E-06 |
| mmu04962 Vasopressin-regulated water reabsorption | 5.38E-07 | 4.78E-06 |
| mmu05222 Small cell lung cancer | 5.92E-07 | 4.87E-06 |
| mmu05200 Pathways in cancer | 5.97E-07 | 4.87E-06 |
| mmu00010 Glycolysis / Gluconeogenesis | 7.46E-07 | 5.85E-06 |
| mmu03420 Nucleotide excision repair | 1.09E-06 | 8.25E-06 |
| mmu03008 Ribosome biogenesis in eukaryotes | 1.23E-06 | 8.95E-06 |
| mmu04910 Insulin signaling pathway | 1.29E-06 | 9.07E-06 |
| mmu04114 Oocyte meiosis | 1.45E-06 | 9.89E-06 |
| mmu05211 Renal cell carcinoma | 1.62E-06 | 1.06E-05 |
| mmu00970 Aminoacyl-tRNA biosynthesis | 1.71E-06 | 1.09E-05 |
| mmu04914 Progesterone-mediated oocyte maturation | 2.06E-06 | 1.27E-05 |
| mmu03018 RNA degradation | 2.92E-06 | 1.75E-05 |
| mmu00520 Amino sugar and nucleotide sugar metabolism | 4.04E-06 | 2.35E-05 |
| mmu00280 Valine, leucine and isoleucine degradation | 6.88E-06 | 3.90E-05 |
| mmu03030 DNA replication | 7.25E-06 | 3.97E-05 |
| mmu00510 N-Glycan biosynthesis | 7.40E-06 | 3.97E-05 |
| mmu00230 Purine metabolism | 8.46E-06 | 4.43E-05 |
| mmu04370 VEGF signaling pathway | 1.71E-05 | 8.71E-05 |
| mmu03020 RNA polymerase | 1.76E-05 | 8.77E-05 |
| mmu04666 Fc gamma R-mediated phagocytosis | 2.19E-05 | 1.06E-04 |
| mmu05219 Bladder cancer | 2.23E-05 | 1.06E-04 |
| mmu04150 mTOR signaling pathway | 5.06E-05 | 2.35E-04 |
| mmu00620 Pyruvate metabolism | 5.92E-05 | 2.68E-04 |
| mmu05214 Glioma | 8.32E-05 | 3.69E-04 |
| mmu04512 ECM-receptor interaction | 1.14E-04 | 4.94E-04 |
| mmu00071 Fatty acid metabolism | 2.08E-04 | 8.82E-04 |

continued on next page

Table S3. KEGG Pathways with Downregulated Genes in Dormant Single Cells, continued

| Pathway Name | p value | q value |
|---|---|---|
| mmu04115 p53 signaling pathway | 2.13E-04 | 8.87E-04 |
| mmu04912 GnRH signaling pathway | 2.28E-04 | 9.31E-04 |
| mmu05210 Colorectal cancer | 2.50E-04 | 9.98E-04 |
| mmu05215 Prostate cancer | 2.68E-04 | 1.05E-03 |
| mmu04722 Neurotrophin signaling pathway | 2.95E-04 | 1.14E-03 |
| mmu04012 ErbB signaling pathway | 3.01E-04 | 1.14E-03 |
| mmu04810 Regulation of actin cytoskeleton | 3.32E-04 | 1.21E-03 |
| mmu00052 Galactose metabolism | 3.33E-04 | 1.21E-03 |
| mmu05223 Non-small cell lung cancer | 3.69E-04 | 1.32E-03 |
| mmu05145 Toxoplasmosis | 4.36E-04 | 1.53E-03 |
| mmu03430 Mismatch repair | 9.59E-04 | 3.31E-03 |
| mmu05213 Endometrial cancer | 1.02E-03 | 3.48E-03 |
| mmu04540 Gap junction | 1.09E-03 | 3.65E-03 |
| mmu05160 Hepatitis C | 1.23E-03 | 4.04E-03 |
| mmu00310 Lysine degradation | 1.93E-03 | 6.26E-03 |
| mmu04210 Apoptosis | 2.24E-03 | 7.11E-03 |
| mmu04966 Collecting duct acid secretion | 2.26E-03 | 7.11E-03 |
| mmu00290 Valine, leucine and isoleucine biosynthesis | 3.31E-03 | 1.02E-02 |
| mmu00630 Glyoxylate and dicarboxylate metabolism | 3.95E-03 | 1.20E-02 |
| mmu04330 Notch signaling pathway | 4.34E-03 | 1.30E-02 |
| mmu04623 Cytosolic DNA-sensing pathway | 4.42E-03 | 1.31E-02 |
| mmu00030 Pentose phosphate pathway | 7.57E-03 | 2.21E-02 |
| mmu04612 Antigen processing and presentation | 8.69E-03 | 2.50E-02 |
| mmu00564 Glycerophospholipid metabolism | 9.59E-03 | 2.71E-02 |
| mmu04664 Fc epsilon RI signaling pathway | 9.69E-03 | 2.71E-02 |
| mmu00051 Fructose and mannose metabolism | 1.02E-02 | 2.82E-02 |
| mmu04621 NOD-like receptor signaling pathway | 1.04E-02 | 2.82E-02 |
| mmu04260 Cardiac muscle contraction | 1.05E-02 | 2.82E-02 |
| mmu05146 Amoebiasis | 1.06E-02 | 2.82E-02 |
| mmu00511 Other glycan degradation | 1.22E-02 | 3.19E-02 |
| mmu00270 Cysteine and methionine metabolism | 1.29E-02 | 3.34E-02 |
| mmu03410 Base excision repair | 1.38E-02 | 3.52E-02 |
| mmu04010 MAPK signaling pathway | 1.47E-02 | 3.70E-02 |
| mmu04380 Osteoclast differentiation | 1.49E-02 | 3.70E-02 |
| mmu03022 Basal transcription factors | 1.76E-02 | 4.32E-02 |
| mmu05020 Prion diseases | 2.00E-02 | 4.86E-02 |
| mmu00250 Alanine, aspartate and glutamate metabolism | 2.04E-02 | 4.89E-02 |
| mmu04964 Proximal tubule bicarbonate reclamation | 2.06E-02 | 4.89E-02 |
| mmu05140 Leishmaniasis | 2.18E-02 | 5.12E-02 |
| mmu00100 Steroid biosynthesis | 2.40E-02 | 5.52E-02 |
| mmu04146 Peroxisome | 2.41E-02 | 5.52E-02 |
| mmu04670 Leukocyte transendothelial migration | 2.58E-02 | 5.84E-02 |
| mmu00480 Glutathione metabolism | 2.76E-02 | 6.18E-02 |
| mmu05218 Melanoma | 2.86E-02 | 6.34E-02 |
| mmu00640 Propanoate metabolism | 3.17E-02 | 6.95E-02 |
| mmu04974 Protein digestion and absorption | 3.38E-02 | 7.34E-02 |
| mmu00900 Terpenoid backbone biosynthesis | 3.42E-02 | 7.35E-02 |
| mmu00330 Arginine and proline metabolism | 3.60E-02 | 7.65E-02 |
| mmu04520 Adherens junction | 3.64E-02 | 7.66E-02 |
| mmu04130 SNARE interactions in vesicular transport | 3.87E-02 | 8.06E-02 |
| mmu04320 Dorso-ventral axis formation | 4.52E-02 | 9.29E-02 |
| mmu05221 Acute myeloid leukemia | 4.56E-02 | 9.29E-02 |

Pathway analysis was completed as described in Methods on single cell samples from the 10X Genomics and the ultralow RNAseq data. Only pathways with q<0.1 are displayed.

**Table S3. KEGG Pathways with Upregulated Genes in Dormant Cells from Ultralow Input Data.** Ultralow RNA-seq analysis was carried out and analyzed as described in Methods. Pathway enrichment was completed using GAGE (see Methods for more detail). Pathways with upregulated genes present are listed with q<0.1.

Table S4. Genes located in differentially open and closed regions of dormant DTCs

| Open Regions | | | | | Closed Regions |
|---|---|---|---|---|---|
| Ccdc54 | BC052040 | Prss48 | Snai2 | Lpar5 | Osm |
| 4930542D17Rik | Aldh8a1 | Atp2c2 | Mybl2 | Pbx1 | Utrn |
| 1700112E06Rik | Rnf216 | Angptl4 | Gtsf1l | Pik3ap1 | Zswim6 |
| E2f3 | Gramd1c | A530016L24Rik | Ankrd6 | | Gorab |
| Klhl5 | Mir3470b | A930011G23Rik | Slc12a8 | | Rin2 |
| Adam19 | Pinx1 | Mpp6 | AI661453 | | Aprt |
| Zbtb40 | Pnkd | Tmem63a | Zfp521 | | Galns |
| 2310008H04Rik | Gm216 | Chp2 | Cdh6 | | Anxa5 |
| C230057M02Rik | Grm5 | Ntf3 | Rgr | | 1810062G17Rik |
| Vars | Frmd4a | Cd209a | Gm6116 | | Grk5 |
| Vwa7 | Vstm4 | A330033J07Rik | Inpp4b | | Sytl3 |
| Sapcd1 | Pde7b | Camk2d | LOC101056207 | | Ezr |
| Msh5 | Fbxl7 | Igsf3 | E2f7 | | Bahcc1 |
| Ninj2 | Gm10523 | Fcrl5 | 1700020G17Rik | | Sc4mol |
| Frmd5 | Nedd4l | Grm7 | Rfx8 | | Plec |
| Gtf2h3 | Gm10743 | Cand1 | Gimap3 | | Ssh1 |
| Tctn2 | Herc3 | 1700054K19Rik | Galnt18 | | Itpkb |
| Scap | Prkch | Srbd1 | Pnpla5 | | Hyal1 |
| Tomm40 | Commd10 | Dcbld1 | Itln1 | | Hyal2 |
| Pvrl2 | Iqgap2 | Eps8 | Hpcal1 | | Hyal3 |
| Sulf1 | 3425401B19Rik | Zfhx3 | Lag3 | | Ifrd2 |
| Pdk4 | St3gal1 | Serpina3g | Ptms | | Nat6 |
| Ankmy2 | LOC101055818 | Serpina3h | A230083G16Rik | | Pgs1 |
| Zdhhc11 | Tiam1 | Hmga2 | Eps8l1 | | Tbc1d10a |
| Anxa5 | Fam65b | 1700006J14Rik | D630041G03Rik | | Timp2 |
| 1810062G17Rik | Galm | 9230105E05Rik | Ppp1r12c | | Arhgef7 |
| Bet3l | 9230110C19Rik | Ctla2a | Tmem141 | | Prkar1a |
| Fam26e | 9530026P05Rik | Tpbpa | Fcna | | Fam20a |
| Bace2 | Gm4211 | 9530026P05Rik | 4933415F23Rik | | |
| Shisa9 | 9330111N05Rik | Masp1 | Rassf1 | | |
| Gm5095 | Coro2b | Epha4 | Gm9917 | | |
| Tie1 | Gstm7 | Acer2 | Tusc2 | | |
| Klf7 | Gstm6 | Sh2d4b | Hyal2 | | |
| Fermt2 | Zfp462 | C330018A13Rik | Hyal1 | | |
| 4930527F14Rik | Plcxd2 | Ccdc60 | Plcb1 | | |
| Atpbd4 | Lrfn2 | Fam188b | Lrba | | |
| Tbcel | Ano6 | Col25a1 | Dpf3 | | |
| Cass4 | Loxhd1 | Itln1 | Akap6 | | |
| Rtfdc1 | Clec14a | Gucy2g | Npas3 | | |
| Gm14047 | Nova2 | Acsl5 | Shc3 | | |
| Krtap13 | Dlc1 | Ccdc22 | Gm4884 | | |
| Efcab10 | Cldn5 | Cacna1f | Cit | | |
| Slc44a1 | Cdc45 | Ear2 | Tnfrsf21 | | |
| Rbms3 | Rapgef5 | Nckap5 | Nlgn1 | | |
| Grm4 | 5830416P10Rik | Atoh8 | Ear1 | | |
| Nrip1 | Fry | Fbxw17 | Slc7a5 | | |
| Rcsd1 | Sfpi1 | Ccser1 | Pdlim3 | | |
| Dapk2 | Mybpc3 | 9130019P16Rik | Gm21190 | | |
| 4931422A03Rik | Gm9292 | Ccdc77 | Ppm1f | | |
| Fbxo3 | Elp3 | Pramef6 | 4933415F23Rik | | |

Genes located in regions of differentially open or closed chromatin, whose $\log_2$fold change≥2 with FDR<0.1 in dormant DTCs.

**Table S4. KEGG Pathways with Downregulated Genes in Dormant Single Cells.** 10X genomics and ultralow RNA-seq analysis was carried out and analyzed as described in Methods. Pathway enrichment was completed using GAGE (see Methods for more detail). Pathways with downregulated genes present are listed with q<0.1.

**Supplemental Materials and Methods**

**MRI and IVIS imaging.** Mice with a DFI>300 days were sent to the Rutgers Molecular Imaging Center and MR images acquired with a 1T M2-High Performance MRI System (Aspect Magnet Technologies Ltd, Netanya, Israel). Mice that had been orthotopically injected with Ink4a cells were injected with 200 µL 15 mg/mL luciferin (Pierce) and imaged using the IVIS Spectrum. LivingImage software was used to set the minimum and maximum luminescent rate to be equal in all images.

**H&E, Masson's Trichrome Staining, and Immunohistochemistry.** H&E staining was completed using xylene for deparaffinization and reducing percentages of ethanol for rehydration. Gill's hematoxylin (Vector Labs) and Eosin-Y Alcoholic stain (Richard-Allan Scientific) was used according to manufacturer's instructions. Slides were dehydrated in increasing percentages of ethanol, placed in 3 washes of xylene for 30 seconds each and mounted using ClearMount (American Mastertech Scientific, Inc). Masson's Trichrome Kit (Sigma) was used according to manufacturer's instructions. Immunohistochemistry was carried out as before (*53*) with E cadherin (Santa Cruz, H108), CD68 (Biolegend, FA-11), and mCherry (Rockland) antibodies.

**DNA/RNA extraction and PCR/qPCR.** DNA was extracted using DNAzol according to manufacturer's instructions. Luciferase presence/absence was measured by using TaqMan gene expression assay kit and Taqman primers (luciferase, Life Technologies/Applied BioSystems) on a StepOne RT-PCR machine (Applied Biosystems). Sequence of forward mutant mouse Kras G12D primer: 5'- ACTTGTGGTGGTTGGAGCAG<u>A</u>-3' and WT Kras forward primer: 5'-ACTTGTGGTGGTTGGAGCTGG-3'. The reverse primer for both primers is 5'-CGTAGGGTCATACTCATCCACA-3'. Standard PCR was carried out using 35 cycles of 95°C for 1 minute, 58°C for 30 seconds and 72°C for 20 seconds. PCR products were visualized on a 2% TAE agarose gel stained with ethidium bromide. RNA was extracted using RNeasy kit (Qiagen) for all standard RNA seq samples or by the Quick RNA miniprep kit (Zymo Research). cDNA was made using the Reverse Transcription Kit (Applied Biosystems) according to manufacturer's instructions. For Kras expression, Power SYBR green (ABI Systems) and mutant G12D Kras and WT Kras specific primers were used. The gene expression level was normalized with β-actin and the average presented with standard deviation.

**ATAC-Seq analysis.** Cells were harvested and frozen in culture media containing 5% DMSO. Frozen cells were sent to Active Motif to perform the ATAC-seq assay. The cells were then thawed in a 37$^{o}$C water bath, pelleted, washed with cold PBS, and tagmented as previously described (*20*) with some modifications (*54*). Briefly, cell pellets were resuspended in lysis buffer, pelleted, and tagmented using the enzyme and buffer provided in the Nextera Library Prep Kit (Illumina). Tagmented DNA was then purified using the MinElute PCR purification kit (Qiagen), amplified with 10 cycles of PCR, and purified. Resulting material was quantified using the KAPA Library Quantification Kit for Illumina platforms (KAPA Biosystems), and sequenced with PE42 sequencing on the NextSeq 500 sequencer (Illumina).

Analysis of ATAC-seq data was very similar to the analysis of ChIP-Seq data. Reads were aligned to the human genome (hg38) using the BWA algorithm (MEME mode; default settings). Duplicate reads were removed, only reads mapping as matched pairs and only uniquely mapped reads (mapping quality >= 1) were used for further analysis. Alignments were extended in silico at their 3'-ends to a length of 200 bp and assigned to 32-nt bins along the genome. The resulting histograms (genomic "signal maps") were stored in bigWig files. Peaks were identified using the MACS 2.1.0 algorithm at a cutoff of p-value 1e-7, without control file, and with the –nomodel option. Peaks that were on the ENCODE blacklist of known false ChIP-Seq peaks were removed. Signal maps and peak locations were used as input data to Active Motifs proprietary analysis program, which creates Excel tables containing detailed information on sample comparison, peak metrics, peak locations and gene annotations. Data tracks were loaded on the Integrated Genome Browser (Bioviz.org) to visualize chromatin open peaks.

**Cell Lines.** The mouse primary pancreatic tumor cell line Ink4a.1 luc/mcherry (Ink4a) was a gift from Dr. Eric Collisson (Stanford University, CA). The metastatic early recurrent Met38 cell line was derived in house from a liver metastasis from a resected mouse. Metastatic late recurrent cell lines 208 and 226-1 were derived *in vitro* by first sorting mCherry$^+$ cells from two separate dormant mice and plating them into a 100 $cm^2$ dish. After feeding them twice a week for over 4 months, colonies formed and were picked to establish the cell lines. All cell lines were confirmed as deriving from the original Ink4a cell line by luciferase absence/presence reaction (data not shown). All lines tested negative for mycoplasma.

**Transfection and Immunofluorescence.** Ink4a.1 cells were plated onto glass coverslips and the next day transfected with the expression plasmid pCMV6-mouse Bhlhe41-DDK-myc (Origene). Two days later, cells were fixed with 4% paraformaldehyde at room temperature for 20 minutes and permeabilized with 0.25% Triton for 10 minutes. Cells were incubated overnight with antibodies to DDK (Origene) and Ki67 (Cell Signaling). Anti-mouse AF488 was used to detect Dec2 (DDK) and anti-rabbit AF594 was used for Ki67. Coverslips were mounted using Vectashield plus DAPI (Vectorlabs) and images taken using the Keyence BZ-X710 fluorescent microscope.

**Flow Cytometry.** One million mouse pancreatic cancer cells from primary tumor cell line Ink4a.1, metastatic early recurrent cell line Met38, metastatic *in vitro* late recurrent cell line 226-1, metastatic *in vitro* late recurrent cell line 208 were used in the Aldefluor assay (StemCell Technologies) according to manufacturer's instructions then stained with PE-Cy5-CD24 (Biolegend), AF700-CD44 (Biolegend), APC-EpCAM (Biolegend), PE-Cy7-CD133 (Biolegend), and PE-Met (EBiosciences), and APC-Cy7-Ki67 (Ki67 from Cell Signaling conjugated to APC-Cy7 using Abcam conjugation kit). For dormant DTCs, minced tissue samples from the liver, brain, heart, intestine, kidney, and lung of dormant (DFI>120 days) mice were placed in DMEM/F12+2% FBS+1% P/S with 200 units/mL collagenase type I, 60 units/mL hyaluronidase, and 50 $\mu$g/mL DNAse I and digested at 37$^o$C for 30 minutes, with vortexing every 5 minutes. Tissue pieces were mashed through a 100 μm cell strainer and spun at 300 x g for 5 minutes. Red blood cells were lysed using ACK lysis buffer for 5 minutes, and then washed once with PBS. Cells were counted and either used for analysis of DTCs for mCherry expression using the BD Biosciences Influx High Speed Cell Sorter or subjected to aldefluor and stem cell marker staining as before. For bone marrow aspirates, femurs of mice were cut on the ends and bone marrow irrigated using PBS into a microcentrifuge tube. Cells were spun at 300 x g for 5 minutes, washed once with PBS, and then subjected to flow cytometry for mCherry expression. For flow cytometry of livers at time of pancreatectomy, mice were injected as described above to create a primary tumor, then 4 weeks later liver was harvested as before. The AbC total compensation bead kit was used according to the manufacturer's instructions to determine compensation for multicolor flow cytometry. For Dec2 staining, Dec2 antibody (Abcam) was conjugated to PE using the PE-conjugation kit (Abcam) according to the manufacturer's instructions. One million cells from dormant mouse livers or from cell lines were stained with 5 μL of PE-Dec2 and CTC markers (except for PE-c-Met) and APC-Cy7-Ki67. For p21, p27, and MHCI expression analysis, liver single cell suspensions were stained with p21-AF488 (Cell Signaling), p27-PE (Cell Signaling), and H-2K$^d$/H-2D$^d$ MHC class I-APC (clone 34-1-2S, detects q haplotype of FVB strain, Biolegend) antibodies. DTCs were first classified as CD44$^+$/CD133$^+$/mCherry$^+$, then analyzed for p21, p27, or MHCI expression. To detect intracellular proteins, cells were fixed and permeabilized using the True-Nuclear Transcription Factor Buffer Set (Biolegend) according to manufacturer's instructions.

**Single cell sequencing data analysis:** Single cells from primary tumor #324, disseminated tumor cells from the liver of dormant mice #239 and #241, and recurrent clone #226 were profiled using 10X genomics platform as per manufacturer's protocol. Approximately 110-120 unique molecular identifiers (UMIs) were identified per sample during single cell RNA-seq profiling. UMIs tagging potential doublets were excluded. We used Seurat package for single cell RNA-seq analysis using published approaches (*38*) (*55*). This pipeline allowed us to conservatively exclude the UMIs with unexpectedly low and high number of expressed genes, a majority of which are likely technical artifacts. Samples #239, #241, #226, and #324 had 95, 95, 71, and 87 unique UMIs (single cell equivalent) in the processed dataset. Approximately 2K-5K genes had detectable expression per UMI (or per single cell), and ~15K -17K genes had detectable expression when reads aggregated over the UMIs were used to estimate sample-level expression. Aggregated sample-level expression estimated from 10X data was significantly correlated with bulk RNA-seq data from biological replicates, indicating that systematic bias is likely low, as reported elsewhere (*56*). The samples were comparable in terms of the number of genes expressed per cell, total number of reads, and other QC measures, and showed no major batch effects. We applied normalization steps, as outlined in Seurat, to compare inter-population single cell expression profiles. We analyzed intra-sample transcriptomic heterogeneity using tSNE and PCA plots. We also performed canonical correlation analysis (CCA), and overlaid expression of selected genes on the tSNE plot to assess specific patterns of transcriptomic heterogeneity.

We created a transcriptomic signature of disseminated tumor cells to identify a set of biomarker genes that are significantly differentially expressed in disseminated cells relative to primary tumor and metastatic lesion consistently in the bulk RNA-seq and 10X genomic data (p-value < 0.05 and fold change >2) using an approach similar to that published elsewhere (*57*).

Raw data from the patient samples were processed following the Drop-seq Core Computational Protocol v2.0.0 from the McCarroll laboratory with default parameters. Briefly, barcodes with low quality bases were filtered out, the resulting transcripts were aligned to GRCH37 using the splice-aware STAR aligner (*58*), and gene-level counts and cell-containing barcodes were estimated. Downstream analyses were performed using the Seurat package (*38*) (*55*). Gene-sets between the enriched liver (NAT) and primary tumor samples were first standardized to include all measured genes in both samples. The NAT and tumor samples were then independently normalized using the sctransform function (*59*) and integrated according to the standard pipeline with the following parameter modifications: for P3552, k.filter=5, and for P7180, k.filter=10, k.score=10, and k.weight=10. We then performed principle component analysis (PCA) with 30 dimensions on the integrated datasets and clustered the cells with default parameters. The cell populations consisting of primary and disseminated tumor cells were identified by comparing their gene signatures with profiles found in scRNAseqDB (*60*), a database of gene expression profiles from annotated single cell RNA-seq experiments. The results were visualized on a t-distributed stochastic neighbor embedding (t-SNE) constructed from the principle components. Overall, we analyzed 409 cells from P3552, of which we identified 22 disseminated tumor cells (DTC) and 47 primary tumor cells, and we analyzed 284 cells from P7180 (58 DTC and 44 primary tumor).

**Tabula Muris Analysis.** We obtained raw read count-based expression data from the Tabula Muris, a compendium of single cell transcriptome data from the model organism Mus musculus, containing nearly 100,000 cells from 20 organs (*21*). We processed and analyzed per gene expression estimates (TPM) from primary, recurrent, and disseminated cells jointly with that from healthy mouse liver, spleen, and pancreas at single cell resolution. The quality control filters implemented in the above analysis were able to identify and exclude UMIs with unexpectedly low and high number of expressed genes in normal tissues, and the results were consistent when tumor and normal samples were processed simultaneously. Normal liver, pancreas, and spleen had 972, 1943, and 1714 unique UMIs (single cell equivalent) in the processed dataset. As before, approximately 2K-7K genes had detectable expression per UMI, and ~15K -17K genes had detectable expression when aggregated to assess sample-level expression. Summary statistics such as the number of genes expressed, quantile of expression estimates, proportion of cells with abnormally high or low expression - were comparable between samples from the tumor and normal tissue sets, and we observed no significant difference between tumor and normal tissue sets in terms of these technical attributes, suggesting that potential batch-effects were reasonably accounted for. We jointly analyzed transcriptomic heterogeneity within and between cell types using tSNE and principal component analysis, as above.

**Human Gene Signature Survival Analysis.** We analyzed gene expression data for the pancreatic cancer patients with short term (median survival 0.8 years, n=15) and long-term survival (median survival not yet reached, n=15) after surgery, and assessed whether the dormancy signature genes show differential pattern of expression between the two groups. We compared gene-by-gene expression between the two groups using the limma R package (*61*), and identified those that show significant between-group differences. We then computed median expression for each gene in each respective group and compared the median expression for the dormancy signature genes between the groups of patients with short- and long-term survival. The gene sets up- and down-regulated in dormancy were analyzed separately. Not all of the genes showed expression data from the dormancy signature. This was due to mouse specific-orthologs and an absence of that specific gene probe on the microarray.

**Methylation-Specific PCR.** Frozen tumor samples were homogenized in Trizol (Sigma). Genomic tumor DNA was then prepared using a Wizard kit (Promega). Dormant $CD44^+$/$CD133^+$/$mCherry^+$ DTCs were sorted by flow cytometry and genomic DNA extracted using DNAzol (Sigma). Where indicated, genomic DNA was methylated with CpG Methyltransferase (M.SssI) (New England Biolabs) as a positive control. For promoter methylation analysis in the CpG island of *Kras*, bisulfite treatment was performed using an EpiMark kit (New England Biolabs). Methylation-specific or unmethylation specific PCR was performed for 50 cycles using Platinum Taq (Thermofisher). PCR annealing temperature was 52 degrees. PCR product size was 163 nucleotides. Methylation-specific primers were: 5'-TAGAATAGGTGGTTTACGTTTTGC-3' and 5'-CTACCTACGCTCCACTCGAA-3'. Unmethylated-specific primers were 5'-GGTAGAATAGGTGGTTTATGTTTTGT-3' and 5'-GAACTACCTACACTCCACTCAAA-3'. For analysis of allele-specific gene body methylation, a gentler bisulfite treatment had to be completed using the Imprint DNA modification Kit (Sigma-Aldrich). PCRs were performed with Platinum Taq (Thermofisher) at an annealing temperature of 50 degrees. Primer sequences for analysis of the gene body G12D methylated allele were 5'-GGATAGTTGTCGATATAATTTCG-3' and 5'-AAATTAACTATATCGTCAAAACGCT-3'. Primer sequences for analysis of the wildtype methylated allele were 5'-GGATAGTTGTTGATAAGTTTATGC-3' and 5'-AAATTAACTATATCGTCAAAACGCT-3'. PCR products were 490 bp in length.

**Gemcitabine Treatment of Dormant DTCs.** DTCs were harvested from livers of dormant mice and prepared for cell sorting as before. Single cell suspensions were stained with AF700-CD44 (Biolegend) and PE-Cy7-CD133 (Biolegend) and sorted based on $CD44^+$/$CD133^+$/$mcherry^+$ expression and allowed to recover for one day in DMEM+10% FBS+1% penicillin/streptomycin. Ink4a.1 and sorted DTCs were treated with 100 nM gemcitabine (obtained through the Rutgers Cancer Institute of New Jersey pharmacy) for 48 hours and cell death assessed using Annexin V and 7-AAD (Guava Nexin reagent, Luminex) staining. Cells were analyzed on the Guava EasyCyte HT flow cytometer (Luminex) in triplicate.
